# Planktonic functional diversity changes in synchrony with lake ecosystem state

**DOI:** 10.1101/2022.06.07.495076

**Authors:** Duncan A. O’Brien, Gideon Gal, Stephen J. Thackeray, Shin-ichiro S. Matsuzaki, Julia L. Blanchard, Christopher F. Clements

## Abstract

Managing ecosystems to effectively preserve function and services requires reliable tools that can infer changes in the stability and dynamics of a system. Conceptually, functional diversity (FD) appears a viable monitoring metric due to its mechanistic influence on ecological processes, but it is unclear whether changes in FD occur prior to state responses or vice versa. We examine the lagged relationship between planktonic FD and abundance-based metrics of system state (e.g. biomass) across five highly monitored lake communities using both correlation and non-linear causality approaches. Overall, phytoplankton and zooplankton FD display synchrony with lake state but each lake is idiosyncratic in the strength of relationship. It is therefore unlikely that changes in plankton FD are identifiable before changes in more easily collected abundance metrics. This suggests that FD is unlikely to be a viable early indicator, but has value as an alternative state measure if considered at the lake level.

**Graphical Abstract:** 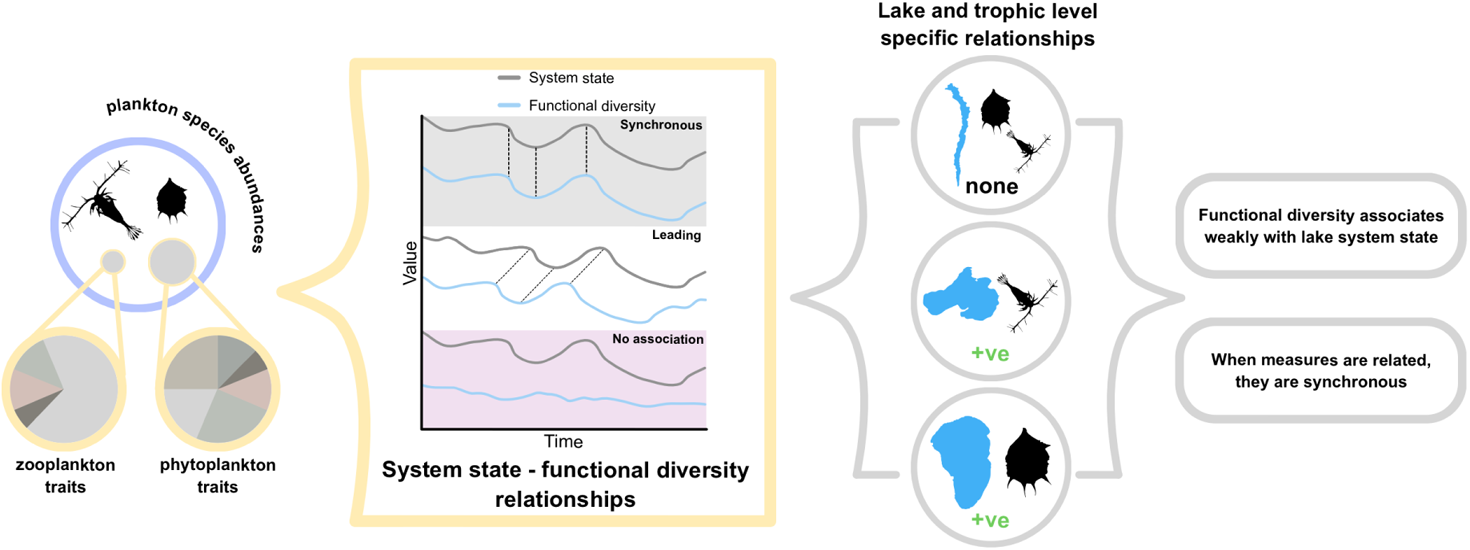

**Data accessibility statement:** Lake Kinneret and Lake Kasumigaura data are available on request, with all other data publicly available and referenced throughout. All code for analysis is available in the Zenodo record (to be released) and the associated GitHub repository (https://github.com/duncanobrien/plankton-FD).

## Introduction

Predicting oncoming ecosystem change is a vital first step in the management of both ecosystems and their associated resources. Ecosystem state or functioning is considered the critical target in this regard as its disruption can result in the loss of many ecosystem services upon which human societies are dependent (Rockström *et al*. 2009). Since the turn of the millennium, biodiversity has appeared as the principal determinant and indicator of both ecosystem state (Tilman *et al*. 2014) and resilience (Oliver *et al*. 2015) across a range of biomes and scales. Biodiversity itself is inherently multidimensional, consisting of taxonomic (i.e. species), functional (i.e. trait) and genetic components amongst others (Lyashevska & Farnsworth 2012; Naeem *et al*. 2016). Despite this multidimensionality, focus has primarily been on the impact of the phylogenetic component with consistent positive associations between species diversity and ecosystem functioning identified in multiple taxa and environments (Cardinale *et al*. 2006; Duffy *et al*. 2017). However, there is significant evidence that other biodiversity dimensions are as impactful on an ecosystem’s function as species richness, particularly the diversity in traits (Cadotte *et al*. 2011; Mouillot *et al*. 2011.

Trait diversity as a predictor of function stems from the view that, ecologically, a species (or a community) is a collection of phenotypic traits that determines their temporal and spatial impact on their surroundings and each other (McGill *et al*. 2006). Higher trait diversity consequently allows species to co-exist and exploit a wider range of ecological niches (Fukami *et al*. 2005) that in turn increases the number of ecological functions/services performed by the community. Multiple studies support this view and present similarly strong associations between functional diversity and functioning measures (de Bello *et al*. 2010; Mouillot *et al*. 2011) with some further suggesting that functional diversity sufficiently outperforms species diversity measures as a predictor of ecosystem state change (Gagic *et al*. 2015; Abonyi *et al*. 2018b). The emerging evidence in favour of functional diversity suggests that trait change can feasibly occur prior to changes in system state, and may represent a viable early warning of change.

Timing is central to managing ecosystems (Hastings 2016); the optimal moment for ecological intervention varies depending on both disturbance severity and the specific system (Walker *et al*. 2014). Consequently, any monitoring strategy should include measures that provide sufficient ‘warning’ to enable appropriate planning and action. The pre-emptive or anticipatory nature of an indicator is therefore key for managers when selecting from a suite of potential indicators (Dale & Beyeler 2001) and mechanistically, functional diversity may fulfil this consideration. Indeed, inclusion of trait information improves the robustness of ecological model predictions (Regos *et al*. 2019; Williams *et al*. 2021) and other early warning techniques (Clements & Ozgul 2016). However, despite these suggestions and repeated claims that functional diversity changes following dramatic state changes (e.g. land use change - Edwards *et al*. 2014; lake regime shifts - Moi *et al*. 2021), there remains a need to confirm that diversity changes also occur prior to ecosystem state change, as required by indicator selection frameworks (Dale & Beyeler 2001). To address this need, we must explore lagged relationships between functional diversity and ecosystem state to identify whether the former provides sufficient warning for management purposes and, if so, over what time horizon.

Lake environments have provided a peerless model for global change ecology as high-resolution data are available from long-term monitoring programmes for multiple sites around the world (Meinson *et al*. 2015). The plankton abundance data from these programmes are increasingly being supplemented with appropriate trait information, allowing lake communities to be classified and organised into discrete functional groupings within and across trophic levels (Reynolds *et al*. 2002; Kruk *et al*. 2011). However, more recently, there has been a shift towards continuous trait measures such as functional diversity (Abonyi *et al*. 2018a; Moody & Wilkinson 2019; Ye *et al*. 2019) facilitated by the emergence of extensive trait databases (Hébert *et al*. 2016; Rimet & Druart 2018) and guidance for trait-based plankton research (Litchman & Klausmeier 2008; Litchman *et al*. 2013). Functional diversity metrics based upon both literature-average and study-collected values have resultingly supported the predictions of biodiversity-ecosystem functioning theory (Tilman *et al*. 2014) by associating strongly with ecosystem functioning. For example, there is a positive correlation between functional diversity and phytoplankton biomass (Vogt *et al*. 2010) as well as displaying a causal relationship with resource use efficiency (Ye *et al*. 2019), Similarly, zooplankton functional diversity correlates with trophic state (Moody & Wilkinson 2019; Moi *et al*. 2021). However, each of these associations were only considered instantaneously when, in fact, lagged/leading associations may have been stronger. Explicit lagged effects are beginning to be considered more widely in system ecology (Gellner *et al*. 2020; Rastetter *et al*. 2021) and biodiversity research (Essl *et al*. 2015), but have not been considered during empirical biodiversity-functioning relationship assessments. Consequently, there are clear knowledge gaps regarding first, whether strong functional diversity-state associations are found consistently over time and among systems, and second, if functional diversity consistently changes prior to changes in commonly used state metrics such as biomass and community composition, allowing it to be a viable and generic leading indicator of ecosystem change.

In this study, we use extensive plankton community datasets from five lakes around the world to assess whether phytoplankton and zooplankton functional diversity changes before, during, or after changes in the state of the lake ecosystems. We quantify the usefulness of functional diversity as a monitoring tool for managers via cross correlation and causation assessments at varying time lags, relative to state. We demonstrate that functional diversity is weakly cross correlated with state, with associations often lake specific and synchronous. Causation assessment via convergent cross mapping yield a similar lack of consistency with bi-directional causal relationships found, implying synchronicity between functional diversity and ecosystem state resulting from stronger extrinsic factors. However, unique dynamics present within the functional diversity time series highlight that due to the multidimensional nature of biodiversity, functional diversity still has value as an alternative measure of state.

## Material and methods

### Lake community data

Lake plankton density (individuals/ml) was compiled from five long term freshwater lake datasets curated by a range of government, university, and not-for-profit sources: Lake Kasumigaura (Takamura & Nakagawa 2012; Takamura *et al*. 2017), Lake Kinneret (Zohary 2004), Lake Mendota (Carpenter *et al*. 2017b, a), Windermere (Thackeray *et al*. 2015) and Lake Zurich (Pomati *et al*. 2020)(Figure S1). As these datasets spanned multiple organisations, countries, and sampling methodologies, we performed a standardisation and quality control workflow. Unidentified and/or unnamed species were removed and if a species was not recording on a sampling date, that species’ density was assumed to be zero. The data was then averaged to mean density per month. To maintain the presence of rare species and better inform functional diversity/community estimates, we only further dropped species if their time series consisted of more than 99% zeroes. A greater presence of zeroes than this prevented the completion of many downstream analyses. Any change in state of these five systems was then quantified from these standardised plankton density data using five metrics, each capturing a different dimension of state change (Table 1).

**Table 1.**
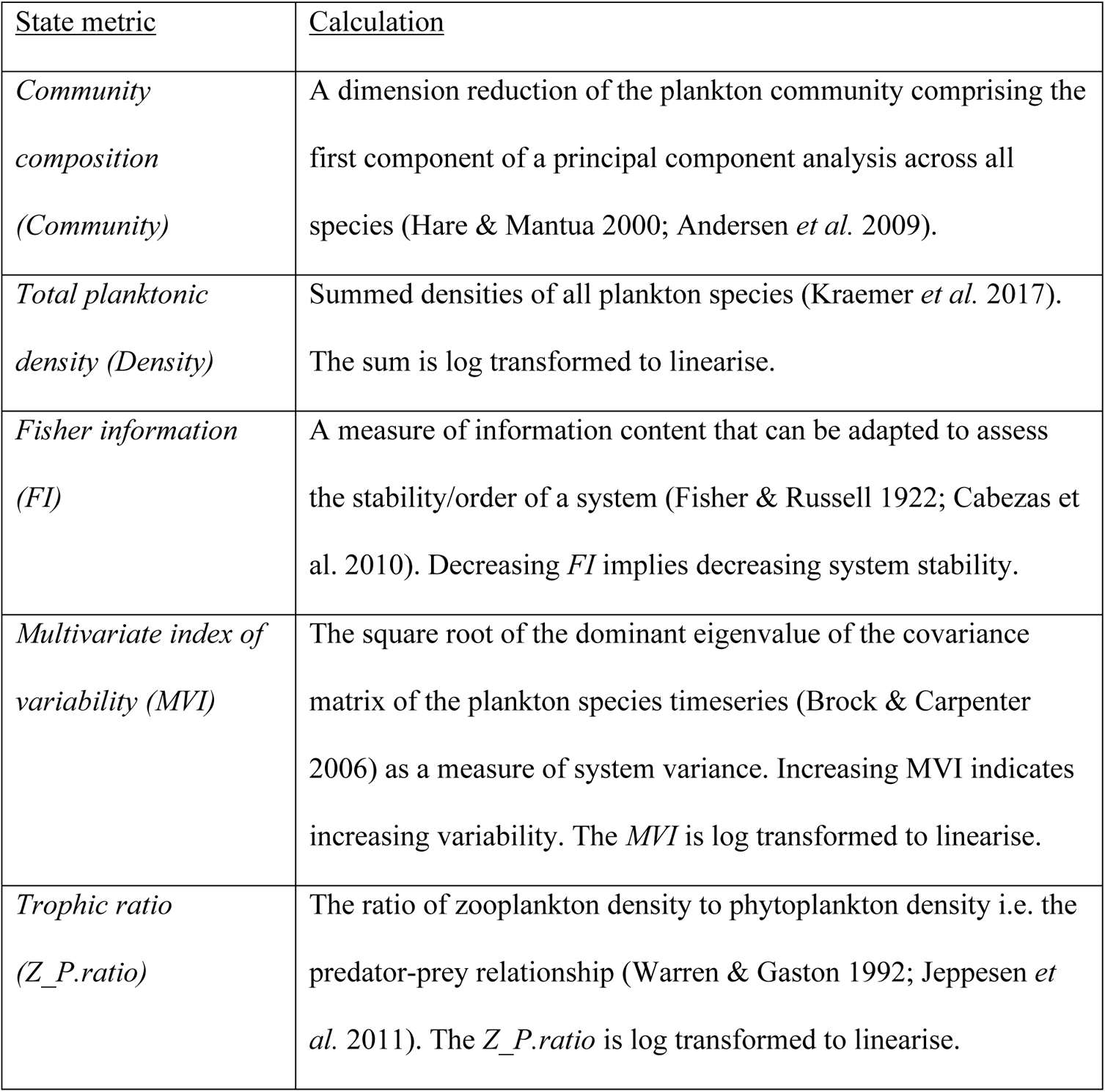
A description of five system state metrics and an exemplary use in the literature

| <u>State metric</u> | <u>Calculation</u> |
| --- | --- |
| *Community composition (Community)* | A dimension reduction of the plankton community comprising the first component of a principal component analysis across all species (Hare & Mantua 2000; Andersen *et al.* 2009). |
| *Total planktonic density (Density)* | Summed densities of all plankton species (Kraemer *et al.* 2017).<br>The sum is log transformed to linearise. |
| *Fisher information (FI)* | A measure of information content that can be adapted to assess the stability/order of a system (Fisher & Russell 1922; Cabezas *et al.* 2010). Decreasing *FI* implies decreasing system stability. |
| *Multivariate index of variability (MVI)* | The square root of the dominant eigenvalue of the covariance matrix of the plankton species timeseries (Brock & Carpenter 2006) as a measure of system variance. Increasing MVI indicates increasing variability. The *MVI* is log transformed to linearise. |
| *Trophic ratio (Z_P.ratio)* | The ratio of zooplankton density to phytoplankton density i.e. the predator-prey relationship (Warren & Gaston 1992; Jeppesen *et al.* 2011). The *Z_P.ratio* is log transformed to linearise. |

### Functional diversity

To underpin the functional diversity estimation, mean species-level trait data were extracted from multiple published databases and articles (Hébert *et al*. 2016; Rimet & Druart 2018; Arcifa *et al*. 2020; Borics *et al*. 2020). Here we consider traits as a measurable characteristic of an individual following Dawson *et al*. (2021). Traits were selected to encompass the three primary ecological axes relevant to phytoplankton (Litchman & Klausmeier 2008; Litchman *et al*. 2013), namely resource acquisition, reproduction, and predator avoidance (Table 2a). Conversely, zooplankton traits are less available and so we followed the suggestions of Barnett *et al*. (2007) and Obertegger *et al*. (2011) to target the same ecological axes (Table 2b).

**Table 2a.**
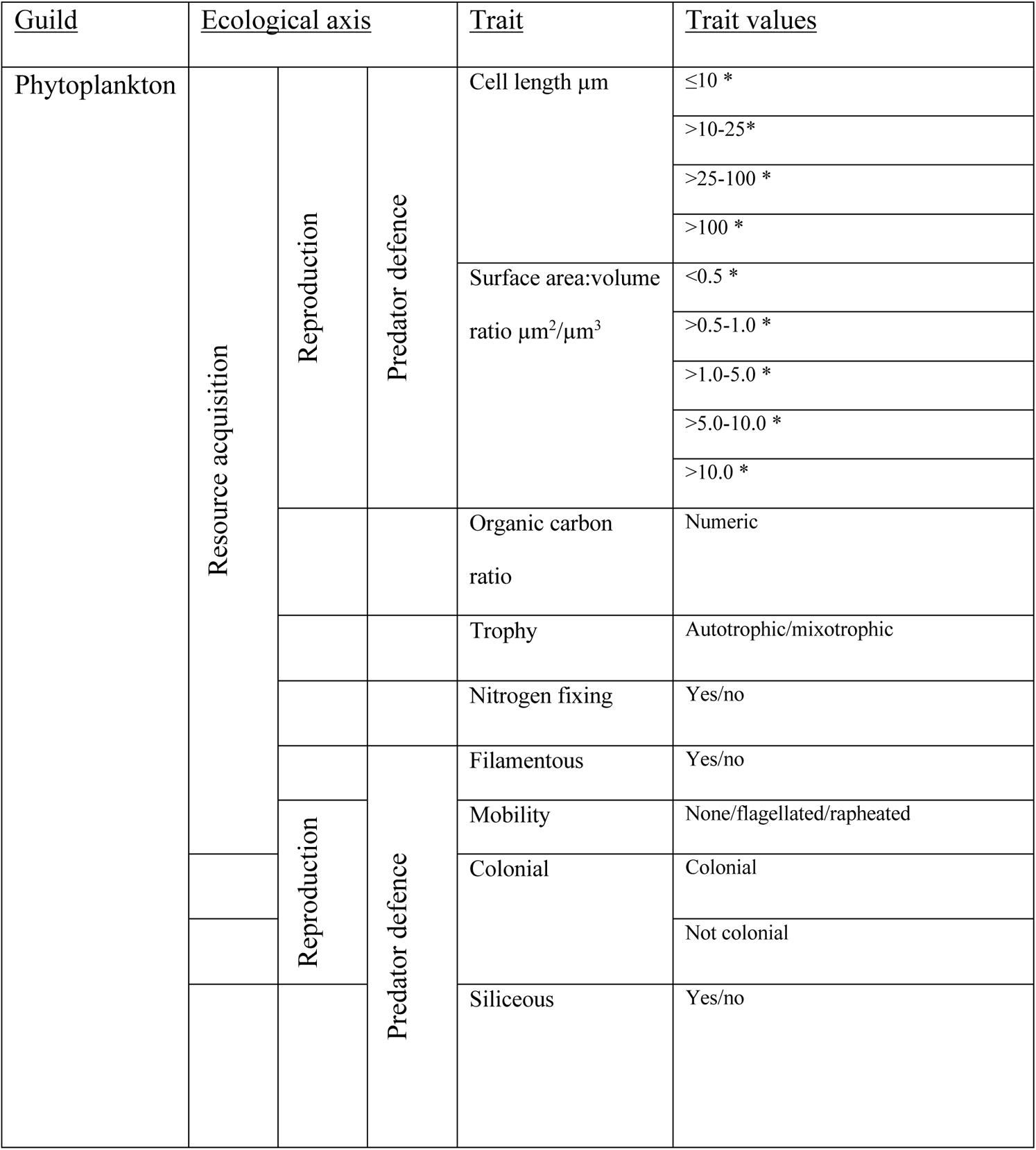
Functional traits of phytoplankton spanning the three primary ecological axes of interest. Quantitative, qualitative, and fuzzily coded values are possible, with fuzzy subcategories indicated by an *.

**Table 2b.**
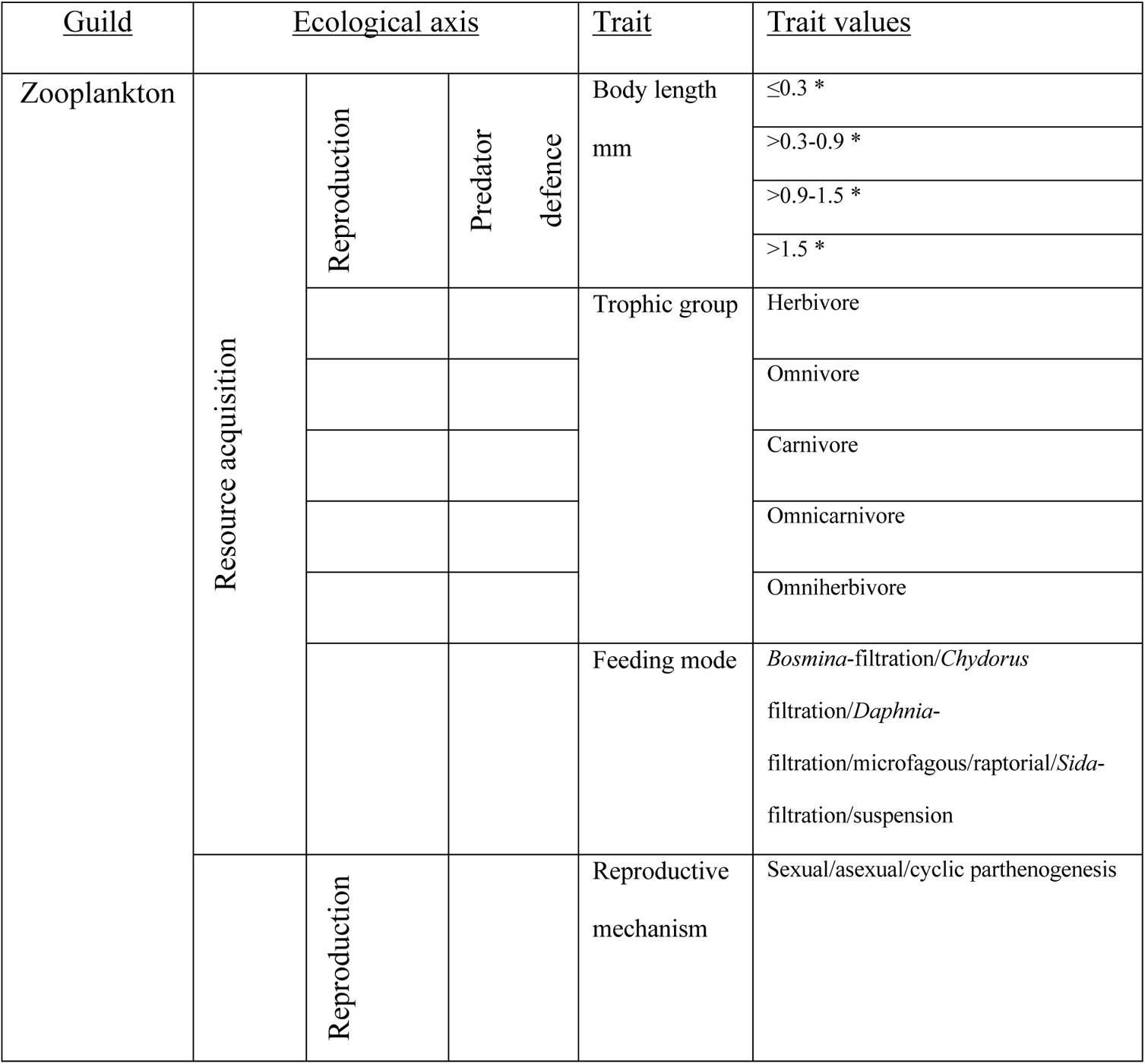
Functional traits of zooplankton spanning the three primary ecological axes of interest. Qualitative and fuzzily coded values are possible, with fuzzy subcategories indicated by an *.

Due to many lake monitoring programs identifying plankton to the genus or family level, it was necessary to integrate multiple species’ trait values into a single taxon value if functional diversity estimation was to be viable. This was achieved via a ‘fuzzy coding’ approach (Chevene *et al*. 1994) which involves the assignment of trait values representing the taxon’s ‘affinity’ to a trait category based upon the variability of species values within it (i.e. the plasticity of the genus). A fuzzily coded trait matrix was therefore uniquely constructed for each lake and plankton guild (phytoplankton vs zooplankton) and from which we calculated a dissimilarity matrix (de Bello et al. 2021 - see Supplementary Information) following the suggestions of Martini *et al*. (2021) for plankton communities. This dissimilarity matrix then underpinned three primary measures of functional diversity (formalised by Pavoine & Bonsall (2011) and Mammola *et al*. (2021)): functional richness (*FRic*), functional dispersion (*FDis*), and functional evenness (*FEve*) (Laliberté & Legendre (2010), Fig. S2). All three measures were computed for each time point using the ‘*mFD’* package (Magneville *et al*. 2022) with a reduced dimension space of 10 to escape generic errors caused during convex hull estimation at higher dimensions. This method therefore results in three functional diversity time series for each lake’s phytoplankton and zooplankton guilds separately.

### Associating system state and functional diversity

Prior to all analyses, system state and functional diversity metrics were scaled to zero mean and unit variance to ensure each shared the same level of magnitude and allow comparison between metrics and lakes. To capture the association between system state and functional diversity, and quantify whether functional diversity leads to changes in state, we performed cross-correlations supplemented by permuted confidence intervals. Each functional diversity measure was cross correlated with each system state metric across a range of lags (from 0 to 60 months), and the observed Pearson correlation coefficient compared to a distribution of pseudorandom correlation coefficients. These coefficients were generated via permutation, where 10,000 surrogate functional diversity time series were constructed from a red/autocorrelated noise process informed by the observed data:

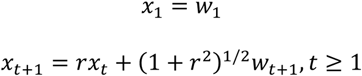

where *r* is the estimated autocorrelation coefficient of the observed time series as estimated by an ARIMA model (Ives *et al*. 2010) and *w* is a white noise process whose mean and variance equalled that of the observed time series. This red noise process consequently generates a surrogate series related to the observed time series but without the sudden changes in trend.

To limit the likelihood of spurious correlation, both the raw time series and permutations were made stationary prior to cross correlation by linear detrending. We then accounted for seasonality by additively decomposing the seasonal component of the original time series (i.e. the mean value for each month, across the length of the time series, standardised to sum to zero) and subtracting this estimate from the model residuals (Zarnowitz & Ozyildirim 2006; Fortin *et al*. 2011).

The observed correlation coefficient at Lag0 and the strongest correlation (positive or negative) across lags was then compared to the permuted 2.5% and 97.5% quartiles to discriminate a stronger cross correlation than expected by serial dependence, and at which lags that may occur. If lags are negative and transgress these ‘confidence intervals’, functional diversity occurs prior to changes in system state, whereas if lags are positive then functional diversity lags state change. Conversely, if the observed correlation resides within the 2.5% and 97.5% quartiles then we consider functional diversity to not correlate significantly with system state.

### Causality and Convergent Cross Mapping

To supplement this cross-correlation approach and provide insight into the information content that functional diversity contains on system state, convergent cross mapping (CCM) was performed on the detrended time series (Sugihara *et al*. 2012). CCM allows the causal influence of one time series on another to be assessed by exploiting a hypothesised shared latent system (Chang et al. 2017; Runge et al. 2019 - see Supplementary Information for details). The presence of forward and reverse causality - functional diversity causing system state and vice versa - for each lake and the optimal time delay (up to Lag60) of causation (Ye *et al*. 2015) was computed using the same permutation method as the cross-correlation approach; if the observed cross map skill (analogous to correlation coefficient) between functional diversity and system state was greater than the 95^th^ quartile of the distribution of cross map skills generated from 10000 surrogate time series, then it was considered significant. Both forward and reverse causality require comparison as, unlike correlation, the strength of relationship depends on the direction of assessment, where strong cross map skills in both directions implies bi-directional causality (Chang et al. 2017). All CCM analysis was performed using the ‘*rEDM*’ package (Park *et al*. 2021).

## Results

### Distinct lake trends through time

We estimated three functional diversity metrics across two plankton trophic guilds in each of five lake monitoring datasets (Figure 1). Lake time series length varied in duration from 24 to 46 years, with a median length of 33 years. The final number of species that contributed to functional diversity estimates varied between lakes due to trait data limitations and longitudinal differences. Consequently, phytoplankton taxa record number ranged from 17 – 130 with a median of 79 taxa, whereas zooplankton records ranged from 4 – 31 taxa with a median of 22.

**Figure 1.**
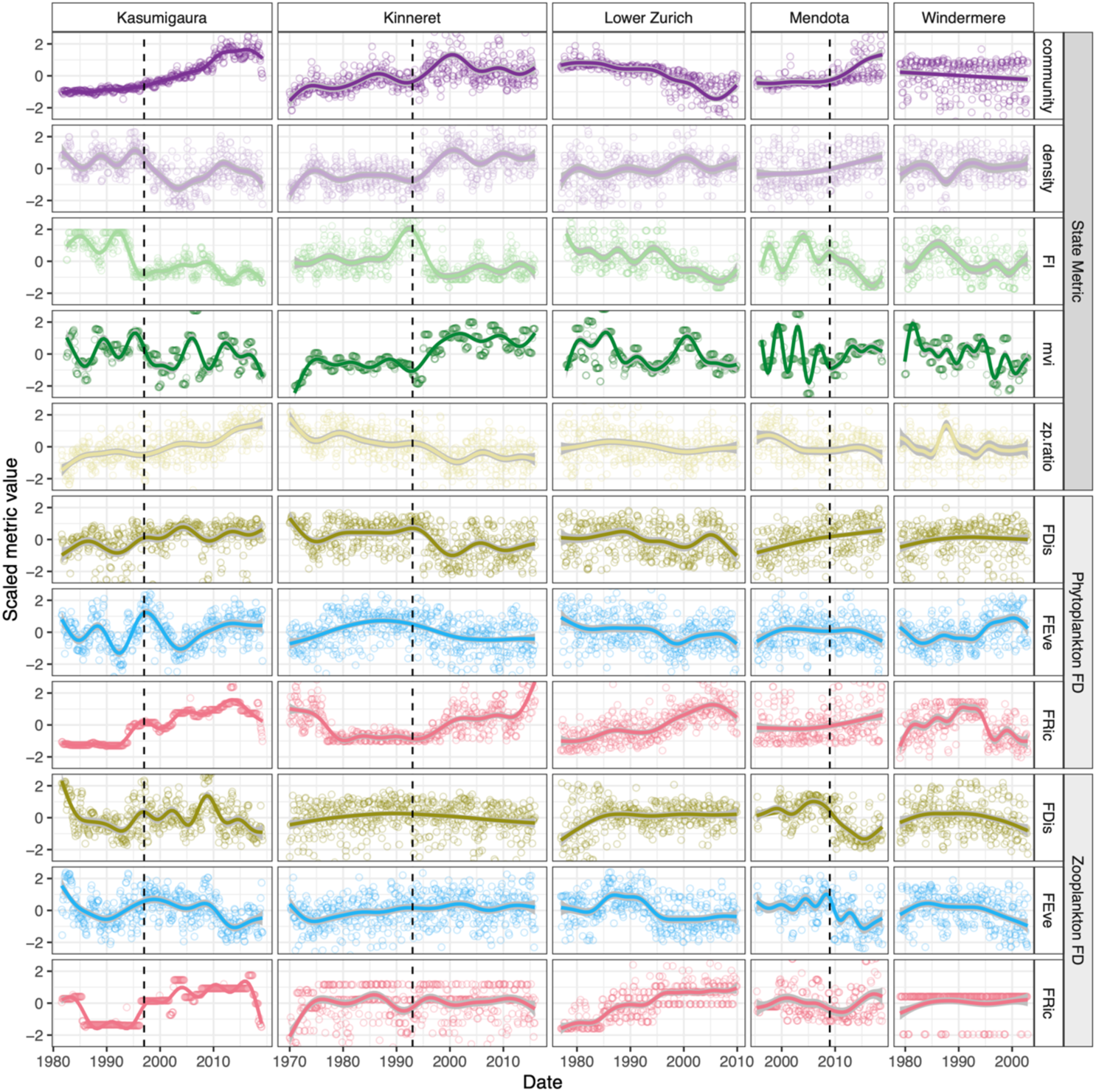
Smoothed time series of the five system state metrics, functional diversity of the two plankton trophic guilds and representation of environmental stressor in each of the five lakes. Smoothed trends are estimated by a generalised additive model of the metric through time and the vertical, dashed line represents literature reported regime shifts. Metric values are scaled to mean zero and unit variance.

All lakes displayed turning points in their system state and functional diversity metrics, implying that over the course of monitoring these systems experienced some form of change in their communities (Figure 1). Most ecosystem state metrics changed simultaneously although Fisher information (*FI*) changed prior to changes in Community, Density and trophic ratio (*Z_P.ratio*), whilst the multivariate index of variability (*MVI*) displayed additional higher frequency fluctuations not identifiable in the other metrics. Functional diversity metrics also displayed unique trends depending on both the system and guild (phytoplankton or zooplankton), for example phytoplankton functional evenness (*phyFEve*) in Lake Kasumigaura displayed abrupt changes compared to other lakes, despite phytoplankton functional dispersion (*phyFDis*) being similar across lakes (Figure 1).

### Synchronicity in both instantaneous and lagged cross-correlation

We consider correlations between functional diversity and system state in the form of both instantaneous/Lag0 correlations (i.e between unlagged time series) and cross-correlations (i.e. when one time series is lagged relative to the other). A strong instantaneous correlation would imply the functional dimension of biodiversity is related to state/functioning, whereas a strong cross-corelation at a negative lag would suggests that functional diversity leads changes in state. However we found correlations were inconsistent across all five of our lake systems (Figure 2, Figure S3), with each combination of functional diversity:system state varying in their proportion of significant correlations (Table S1). The phyFDis:Density relationship expressed the strongest average correlation at Lag0 (median ± se: −0.40 ± 0.06) across all lakes and was significant in four of the five. The only other relationships of a similar magnitude were phyFDis:Z_P.ratio (0.29 ± 0.06) and phyFEve:Density (−0.22 ± 0.06) each of which was also significant in four of the five lakes. Conversely, zero significant correlations were observed between phyFRic:Z_P.ratio, zooFDis:MVI, zooFEve:MVI, zooFRic:Density, zooFRic:MVI and zooFRic:Z_P.ratio. When considered in isolation, Lake Kinneret (which showed the fastest and most distinct change in system state) displayed significant relationships for all but two combinations of indicators and phytoplankton functional diversity, (phyFRic:Community and phyFRic:Z_P.ratio), whereas Windermere (whose state is most stationary) displayed no significant zooplankton relationships. Our results suggest that functional diversity is not universally correlated with system state in lake systems, but rather it is the unique dynamics/context that dictate(s) the strength of the relationship. One possible exception to this is phyFDis, which often strongly correlates with Density and Z_P.ratio.

**Figure 2.**
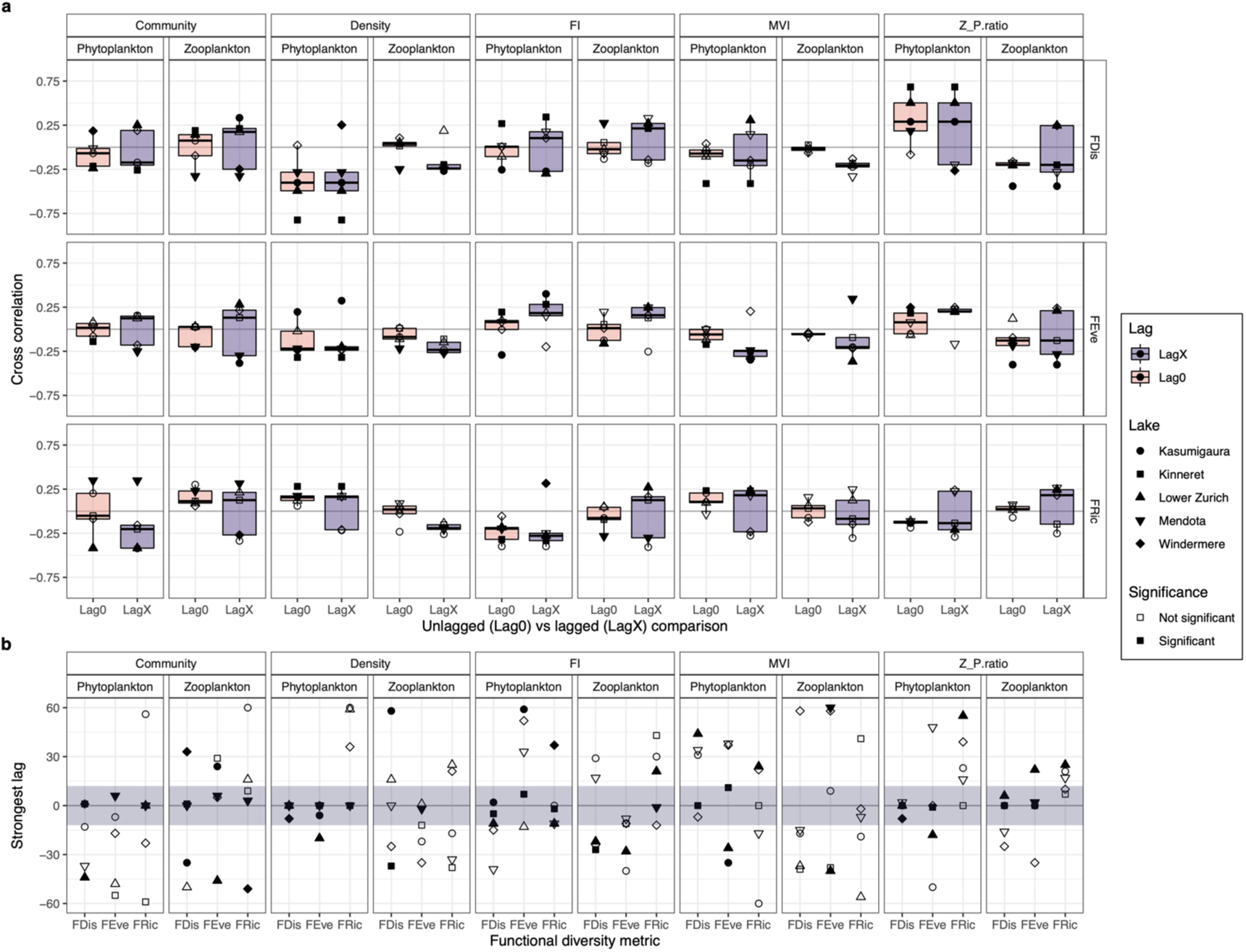
a) Boxplots of cross correlations between each system state and functional diversity metric combination, estimated when functional diversity was unlagged relative to system state (Lag0) versus when it was lagged (LagX). These comparisons have then been stratified by functional diversity metric (FDis, FEve, FRic), state metric (Community, Density, Fisher information, Multivariate variance index, and Trophic ratio) and trophic level (phytoplankton vs zooplankton). A filled point indicates that the mapping was in the strongest 5% of permuted mappings and is considered significant. LagX values represent the strongest cross map skill estimated separately for each lake and across all lags (−60 to +60 months). Consequently, lakes often displayed different strongest lags. b) The spread of those lags across lakes for each functional diversity and system state metric combination. The dark band in panel b represents a ± 1 year lag/lead, which, if a significant (filled) point is found, is considered a synchronous change between the functional diversity and state metric.

When we considered lagged cross-correlations (LagX), most relationships increased in average absolute correlation coefficient (Table S2), but only 13 of these relationships increased their proportion of significant correlations (14 others remained unchanged and three decreased). The phyFDis:Density relationship noted above emerged as universally shared (Figure 2, Figure S4), with phytoplankton functional diversity overall displaying stronger and more consistent relationships with system state than zooplankton functional diversity (with 35 significant relationships compared to zooplankton’s 26 [Figure 2, Figure S4]). The optimal lag differed between the different functional diversity:system state relationships; for example, the phyFDis:Density relationship remaining strongest at lag0 (median correlation ± se: −0.40 ± 0.08, median lag months ± se: 0 ± 0.72), whilst phyFDis:Z_P.ratio (0.29 ± 0.08, 0 ± 0.78), phyFRic:FI (−0.28 ± 0.06, −2 ± 3.98) and phyFEve:MVI (−0.25 ± 0.05, 11 ± 6.86) combinations did appear equivalently strong but cluster within ±12 months of Lag0 (Figure 2d, Figure S5). These results indicate general synchrony between functional diversity and system state if the system expresses any relationship at all. Ultimately, cross-correlations do not reveal clearly general leading nor lagging changes in functional diversity relative to system state but phyFDis is a good covariate of total planktonic density and trophic ratio at Lag0.

### Convergent cross mapping reveals specific relationships of importance

When causal relationships were estimated using convergent cross mapping, many of the observed weak cross-correlations at Lag0 display causal forcing, although the more nuanced approach indicates that many of the previously identified correlations (Figure 2) may be spurious (Figure 3). For example, phyFDis:Density and phyFDis:Z_P.ratio mappings remain significant in all lakes but we found most mappings were within the permuted null distribution (Figure S6), suggesting no causal relationship between functional diversity and system state. The strongest average cross map skills were estimated for phyFRic:MVI (median ± se: 0.37 ± 0.04), zooFDis:MVI (0.26 ± 0.02), phyFEve:MVI (0.24 ± 0.02) and phyFDis:Density (0.23 ± 0.05) although the variation was high between lakes (Figure S6, Table S4). Causality was also often found for the reverse relationship, where functional diversity maps system state, with the majority of the significant relationships expressing bidirectional causality (Figure S7, Table S4, Table S5) – i.e. both functional diversity and system state influence one another, rather than one being a product of change in the other. This does not support strong causal relationships between functional diversity and system state, and suggests that the previously identified synchronous Lag0 correlations result from not the influence of diversity but from other extrinsic factors.

**Figure 3.**
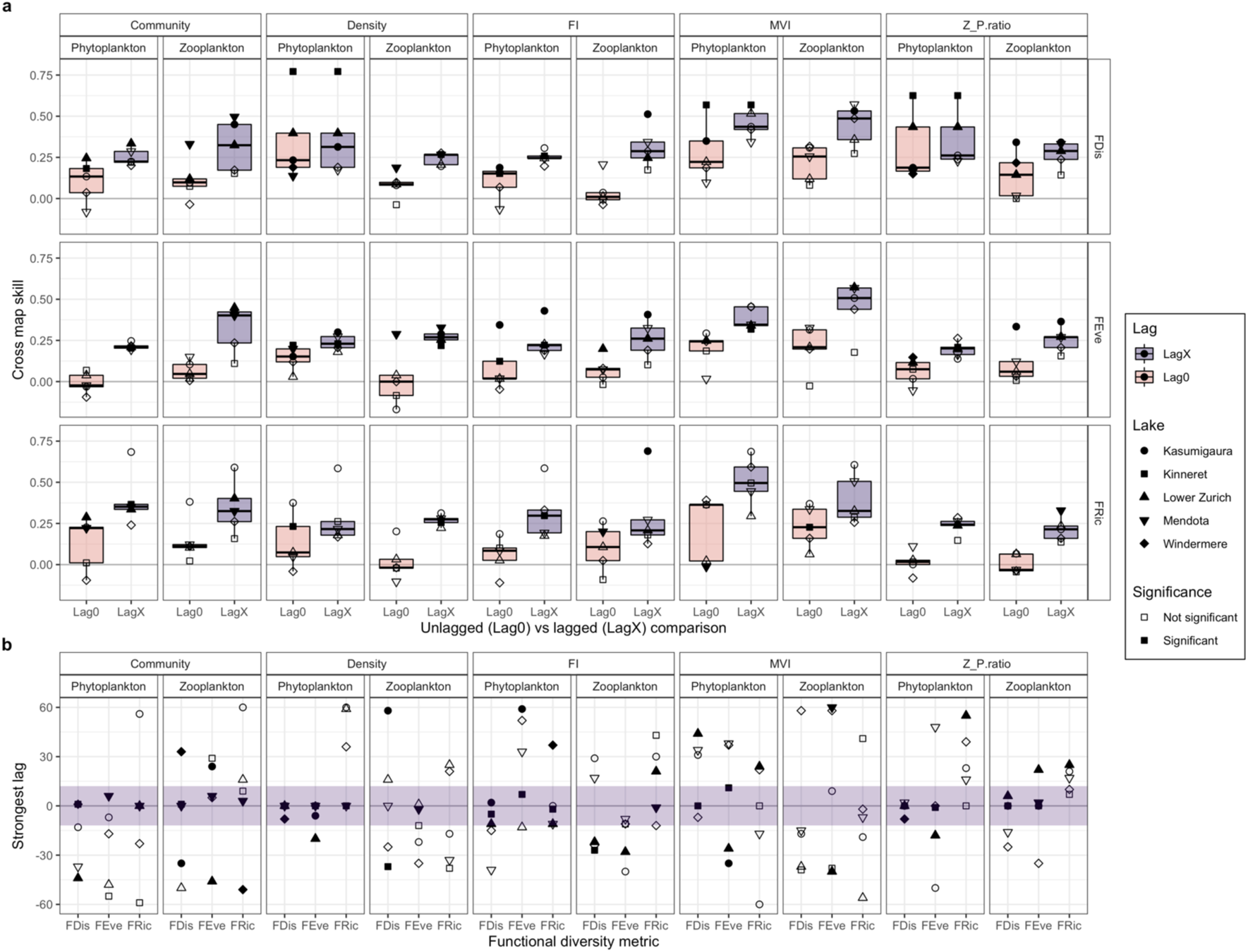
a) Boxplots of cross mapping skills between each system state and functional diversity metric combination, estimated when functional diversity was unlagged relative to system state (Lag0) versus when it was lagged (LagX). These comparisons have then been stratified by functional diversity metric (FDis, FEve, FRic), state metric (Community, Density, Fisher information, Multivariate variance index, and Trophic ratio) and trophic level (phytoplankton vs zooplankton). A filled point indicates that the mapping was in the strongest 5% of permuted mappings and is considered significant. LagX values represent the strongest cross map skill estimated separately for each lake and across all lags (−60 to +60 months). Consequently, lakes often displayed different strongest lags. b) The spread of those lags across lakes for each functional diversity and system state metric combination. The dark band in panel b represents a ± 1 year lag/lead, which, if a significant (filled) point is found, is considered a synchronous change between the functional diversity and state metric.

Interestingly, introducing lags improved the strength of causality by an average of 0.18 skill (implying that stronger causal relationships could be estimated from lagged data), yet did not increase the proportion of causal cross mappings (Figure 3, Figure S8). With lags, no relationship was significant in all lakes with only one significant in four of the five: phyFEve:Density (Figure S8, median skill ± se: 0.28 ± 0.00; median lag months ± se: −51 ± 8.29). Only the phyFRic:Density, phyFRic:MVI, and zooFRic:MVI mappings remained universally non-causal however. The optimal lag also differed between mappings with CCM displaying greater variation than that of the cross-correlations and less prevalence within ±12 months (Figure 3d, Figure S9, Table S6). It is worth noting that counter to the cross-correlation assessment, zooplankton functional diversity had a larger proportion of significant mappings than phytoplankton (37% versus 27%).

Figure 4 presents the relative direction of causality between functional diversity and system state using the strongest lag as a method to disentangle forward causality from bidirectional. Using the zooFDis:Density relationship as an example (Figure 4a, second column), one forward causality assessment (zooFDis ‘causing’ Density) was significantly stronger than the permuted distribution (Kinneret) versus two reverse assessments (Density ‘causing’ zooFDis: Kasumigaura and Lower Zurich), but Kasumigaura’s reverse assessment occurred at positive lags whereas the forward assessment occurred at negative lags. This crossing of the central 0 month line of the dashed pairing line indicates that zooFDis’ information leads Density’s and that the influence of zooFDis synchronises the two measurements. The opposite is true for Lower Zurich, whereas the flat line in Kinneret and Mendota suggest equal influence. If all lakes are considered together, then multiple crossing of pairing lines suggests no consistent causal relationship. Therefore, overall, considering the relative direction of causality between functional diversity and system state irrespective of significance, strong overlaps were identifiable between forward and reverse mappings for the majority of associations (Figure 4, Figure S10, Table S7, Table S8). The exceptions to this trend included FI leading phyFDis (Figure 4a) and both phytoplankton and zooplankton FEve (Figure 4b). The reverse was true with Z_P.ratio being led by all but zooFDis and zooFRic, and Density being led by both phytoplankton and zooplankton FRic (Figure 4c). These results imply functional diversity and system state are not strongly causally related, but both contain equivalent information on each other, with no consistent leading or lagging causality (Table S5, Table S6). Thus, most relationships are synchronous and supports the overall cross-correlation assessment. The FDis and Density relationship is identified as the most robust correlation and cross mapping however, and therefore represents the one strong functional diversity:system state association across lakes and trophic levels.

**Figure 4.**
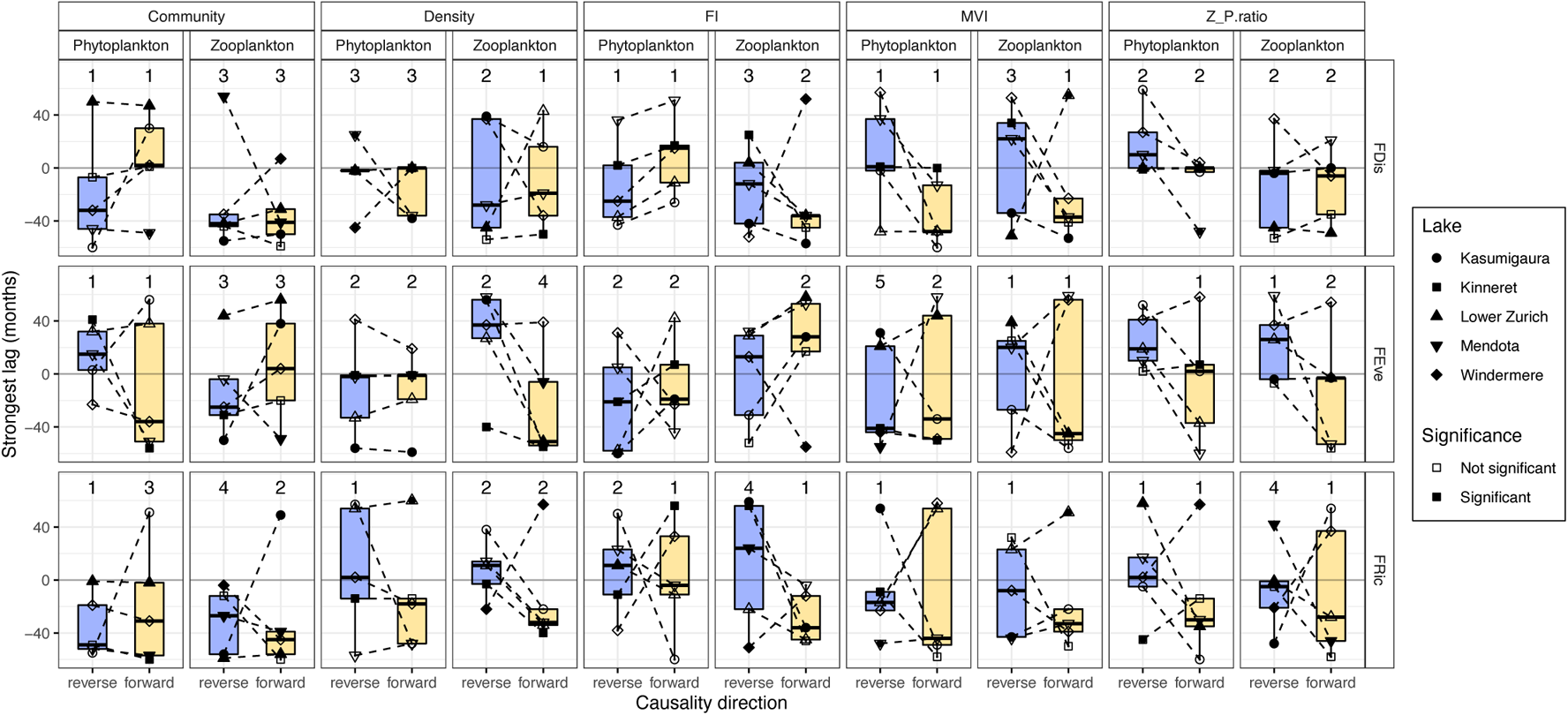
Boxplots of the paired lags between forward and reverse causal estimates for each functional diversity:system state combination. These comparisons have then been stratified by functional diversity metric (FDis, FEve, FRic), state metric (Community, Density, Fisher information, Multivariate variance index, and Trophic ratio) and trophic level (phytoplankton vs zooplankton). Filled points represent a significant causal relationship and the reported value is the number of significant mappings (out of five). Dashed lines link the two paired estimates (forward and reverse mappings within a lake). If one of these pairing lines crosses the grey, central lag line, then one of the metrics has a delayed impact and exerts sufficient causation on the other that synchronicity may occur. Variable directions in line/a flat line across all lakes can be interpreted that both metrics have equivalent causal delays upon each other.

## Discussion

Here, we compared how planktonic functional diversity estimated from literature average trait values changes relative to abundance-based metrics of system state. We find that simple correlative techniques described no or very weak synchronous relationships between functional diversity and state, a finding which is not substantially improved when lags between time series are considered. However, phytoplankton functional dispersion (*phyFDis*) was universally related with total planktonic density (*Density*) albeit synchronously. Conversely, bi-directional causality was relatively prevalent between functional diversity and state when assessed using convergent cross mapping, although Fisher Information (*FI*) often led functional diversity. In both approaches individual lakes expressed unique association strengths which limits our ability to make general conclusions on the use of functional diversity as a warning of ecosystem change. Functional diversity appears not to change consistently prior to abundance-based system state metrics overall but display unique dynamics distinct from state that may indicate their use as an alternative measure rather than a pre-emptive indicator.

The absence of strong relationships was unexpected considering the bulk of literature reconciling ecosystem state with the phylogenetic and functional dimensions of biodiversity, both in planktonic (Abonyi *et al*. 2018b; Moody & Wilkinson 2019; Ye *et al*. 2019) and non-planktonic communities (Dıaz & Cabido 2001; Cadotte *et al*. 2011; Gagic *et al*. 2015). However, it is not a universally identified relationship, with multivariate functional diversity measures failing to predict alpine biomass production (Zhu *et al*. 2016), the strength of relationship varying with disturbance in stream plant communities (Biswas & Mallik 2011), and tree carbon stocks responding uniquely to individual forests’ functional diversity (Ruiz-Jaen & Potvin 2010). Crucially, it is in controlled experiments that strong, consistent relationships are found whereas observational studies are more variable due to abiotic and biotic interactions filtering the taxa present at any moment in time to a ‘realised’ level of biodiversity that differs to the ‘true’/initial diversity examined in experiments (Hagan *et al*. 2021). We identify similar ambiguity here, with distinct relationships for individual lakes, each of which is known to be experiencing different levels of external stress. For example, Lake Kasumigaura and Lake Kinneret are considered to have undergone a regime shift in the late 1990s (Fukushima & Arai 2015) and mid-1990s (Roelke *et al*. 2007) respectively whereas Windermere is relatively stable. Our system state metrics identify those two rapid regime changes and exhibit the strongest associations with functional diversity. Lake Kinneret in particular displays a sudden but relatively brief change in all its state metrics which is mirrored in its phytoplanktonic functional dispersion (*phyFDis)*.

It is likely the rapid change in state is the driver of the observed strong correlations between functional diversity and system state in Kinneret compared to the others, where the magnitude of change enforces an instantaneous shift in all the metrics we explored. This is supported by our paired convergent cross mappings in the circumstances where the lagging variable displays positive lags and the leading variable displays negative lags. When strong forcing is applied to a coupled system, the phenomenon of generalized synchrony can occur (Rulkov *et al*. 1995) as one system component exerts sufficiently strong causation on another that it brings them in to alignment/synchrony. Therefore, while synchronicity may be visible at short time scales, the leading variable is in fact exerting strong causality to synchronise the two (Ye *et al*. 2015). In ecology, the Moran effect (Moran 1953) describes the phenomenon at macrospatial scales, with regime shifts acting as temporal analogues (Wernberg *et al*. 2013). The ubiquitous association in Lake Kinneret matches these examples as the synchrony strengthens for the short periods during the regime transition to improve the overall correlation. This implies that, whilst functional diversity’s relationship is system specific, during regime transitions strong changes can be identified alongside typical system state measures, but does not pre-empt them at management relevant timescales.

The differences between convergent cross mapping (CCM) and cross-correlation in characterising the overall relationship between system state and functional diversity supports the work of Sugihara and colleagues (2012) who show that traditional regression methods are unable to accurately identify complex associations between related ecological time series. Chang *et al*. (2022) also identified chained feedback effects in phytoplankton networks using the technique. Indeed, while non-linear mappings revealed less stronger-than-null relationships than the correlative approach, CCM highlights the insightful and non-spurious relationships of causal value. We find agreement that FDis is linked with total plankton density and that – when causation is present – typically both system state and functional diversity exert equal effects upon each other. In this regard, we consider the two measures as changing together, possibly in response to an unmeasured environmental variable. However, time delay effects are evident for certain metrics, particularly those involving FI. Fisher information has previously been suggested to pre-empt regime shifts in long time series (Cabezas *et al*. 2010; Spanbauer *et al*. 2014; Ahmad *et al*. 2021), where decreasing FI indicates decreasing stability of the system. There has been no extensive assessment of FI’s capability in natural environments but, qualitatively, FI appears to change trajectory prior to each major turn point in the lakes explored in this study and can cross map/‘cause’ change in functional diversity.

One key difference between studies that may limit our identification of strong associations is the length of time series. We find more significant relationships in the longer time series (Lake Kasumigaura, Kinneret and Zurich) than the short (Lake Mendota and Windermere). While this provides more data points for both correlation and CCM, our conservative approach of detrending and referencing an autocorrelated, permuted null distribution mitigates the likelihood of spurious correlations resulting from the shared system and larger datasets (Calude & Longo 2017). As a result, we believe our results are valid.

Similarly, the ability of functional diversity to pre-empt system change may be hampered by the quality of estimates from literature average values. Hutchinson’s paradox (Hutchinson 1961) highlights the high niche overlap of many planktonic species and resulting similarity in many routinely measured traits. This results in a community consisting of many functionally similar species when quantified from traits such as cell length or nitrogen-fixing ability. It was this rationale that led to the development of Reynolds’ phytoplankton functional groupings (Reynolds *et al*. 2002) to circumvent this apparent niche overlap and may indicate a weakness of the continuous functional diversity approach we applied here. While this complication is magnified by the lack of system specific trait information, it is our belief we were able to identify sufficiently distinct diversity trends and relationships using the proposed average-trait framework of Martini *et al*. (2021) to validate the approach.

The use of average trait values is further validated by our understanding that the complex, multi-trophic interactions occurring in these diverse lake communities minimise competitor exclusion and facilitate species with overlapping niches (Brose & Hillebrand 2016; Albert *et al*. 2021). We see differences in the two plankton trophic levels’ diversity but none sufficiently consistent to describe a universal pattern of top-down/bottom-up control. Strength of phytoplankton-zooplankton trophic coupling does vary with the degree of lake oligotrophication (Carney & Elser 1990; Bernat *et al*. 2020; Dong *et al*. 2021) and presumably the lack of consistency results from the variable importance of phytoplankton versus zooplankton guilds in structuring the lake community. Zooplankton is particularly important in Lake Mendota for example, where the appearance of the invasive spiny water flea (*Bythotrephes longimanus*) (Walsh *et al*. 2017) decimated the zooplankton assemblage and initiated a trophic cascade towards a turbid, phytoplankton-dominated community. This importance is identified in both our correlative and causality assessments, with minimal phytoplankton functional diversity associations with Mendota’s system state compared to the improved pre-emptive performance of certain zooplankton metrics. Typology and nutrient status may also be influencing the variable association strength between functional diversity and system state across lakes, but this requires a larger lake dataset to identify. Consequently, when performing functional diversity assessments in lake ecosystems, it appears necessary to consider each system independently within its own context.

The use of lakes for functional diversity research may also avoid previous concerns that our understanding of the mechanistic relationship between biodiversity and ecosystem functioning stems from field experiments where biodiversity effects can only be considered as ‘local’ (Hagan *et al*. 2021; Thompson *et al*. 2021). Island habitats have previously been considered as key study systems to assess the impacts of biodiversity due to their defined taxa pools matching the assumptions of much biodiversity-ecosystem functioning conceptual research (Kardol *et al*. 2018). Lakes may be considered biogeographically insular (MacDonald *et al*. 2018) with dispersal between neighbours restricted compared to the terrestrial environments underpinning much of our understanding of biodiversity-functioning relationships. Thus, the context surrounding lake plankton functional diversity may better represent the theoretical local level effects of biodiversity.

To conclude, the synchronous association between functional diversity and system state conflicts with the conceptual mechanistic relationship between biodiversity and ecosystem functioning. Most likely, any delayed impacts of functional diversity on our selected state measures are insufficiently long to warrant the use of functional diversity as an early indicator of ecosystem change, although the system-specific dynamics of the functional metrics do sometimes yield unique dynamics not seen in the state measures. The relationship between functional diversity and ecosystem state will ultimately depend on the combination of environmental stressors, traits present, and taxa interactions, which together potentially mask the overall relationship. Trait information is still vital to support our understanding of total biodiversity change, but dimensionally reduced trait measures like functional diversity are less informative than practical abundance-based measures (such as Fisher information) to ecosystem managers.

## Acknowledgments

DAO received funding from the GW4+ FRESH Centre for Doctoral Training in Freshwater Biosciences and Sustainability (NE/R011524/1). We thank Tamar Zohary for the long-term phytoplankton dataset from Lake Kinneret, and Heidrun Feuchtmayr for providing data from Windermere; monitoring at this site is currently supported by Natural Environment Research Council award number NE/R016429/1 as part of the UK-SCAPE program delivering National Capability. We also thank the field and laboratory teams who have collected all of the data used in this study. We also declare we have no competing interests.

## Notes

https://github.com/duncanobrien/plankton-FD

### Competing Interest Statement

The authors have declared no competing interest.

