## Supplementary Information for "Planktonic functional diversity changes in synchrony with lake ecosystem state"

Supplementary Methods

*Fuzzily-coded trait dissimilarity*

Fuzzy coding is an approach that has been successfully applied to multiple freshwater macroinvertebrate communities (Brown *et al.* 2018; Múrria *et al.* 2020), and has been suggested as an appropriate tool to circumvent some of the challenges non-species level data can bring to trait-based approaches in plankton (Martini *et al.* 2021). In practice, a fuzzy trait is sub-categorised and ‘affinity’ is the proportion of the taxon’s species which fall within each of the fuzzily coded sub-categories (Gower 1971; de Bello *et al.* 2021). During this study, each of the lakes have sufficient monitoring documentation that genus level data identifies the species that have had their abundances pooled to form a taxon count. Therefore, we were able to code only species known to contribute to the count rather than coding all possible taxon members present in the trait databases. A fuzzily coded trait matrix was consequently uniquely constructed for each lake and plankton guild (phytoplankton vs zooplankton), and from it a dissimilarity matrix based on a modified Gower index was derived using the ‘*gawdis’* package in R (R Core Team 2020). This dissimilarity matrix was then Cailliez transformed (Cailliez 1983) to improve suitability for Euclidean-based analysis during construction of the ‘trait space’. Our choice of a dissimilarity-based functional diversity methodology stems from our aim to quantify changes in trait space size through time (Mammola *et al.* 2021).

*Convergent cross mapping*

Convergent cross mapping invokes Takens’ embedding theorem (Takens 1981) which suggests that an underlying latent system/attractor manifold can be reconstructed from one or more related time series. Specifically, if an observed time series is considered a transformation of the manifold’s states over time, transformed by an observation function consisting of stochasticity and observation error, then the manifold’s reconstruction is possible by time-delay embedding the time series (Chang *et al.* 2017). As a result, if two time series, X and Y, share a manifold, a causal relationship between them can be assessed. Causality is identified by comparing the quality of prediction of one time series from the reconstructed system built upon the time-delay embedding of another (Runge *et al.* 2019); if X can be predicted using the reconstruction from Y, then X had a causal effect on Y. The reciprocal relationship is then also tested to establish whether bi-directional causation is present.

The appropriate embedding dimension for CCM was selected using nearest neighbour forecasting, specifically simplex projection (Sugihara & May 1990). Simplex projections generate forecasts of the manifold reconstruction and identifies the embedding dimension with the highest prediction skill (represented as the correlation between observed and predicted values). In this analysis, we explored dimensions from 0 to 10 with an increment of 1, using *𝜏* = 1 (a month) for embedding. Using the identified embedding dimension, CCM analysis was performed for each time-to-prediction lag/delay with a library size set to the maximum possible to improve prediction error and the likelihood of convergence (Sugihara *et al.* 2012).

Supplementary Figures

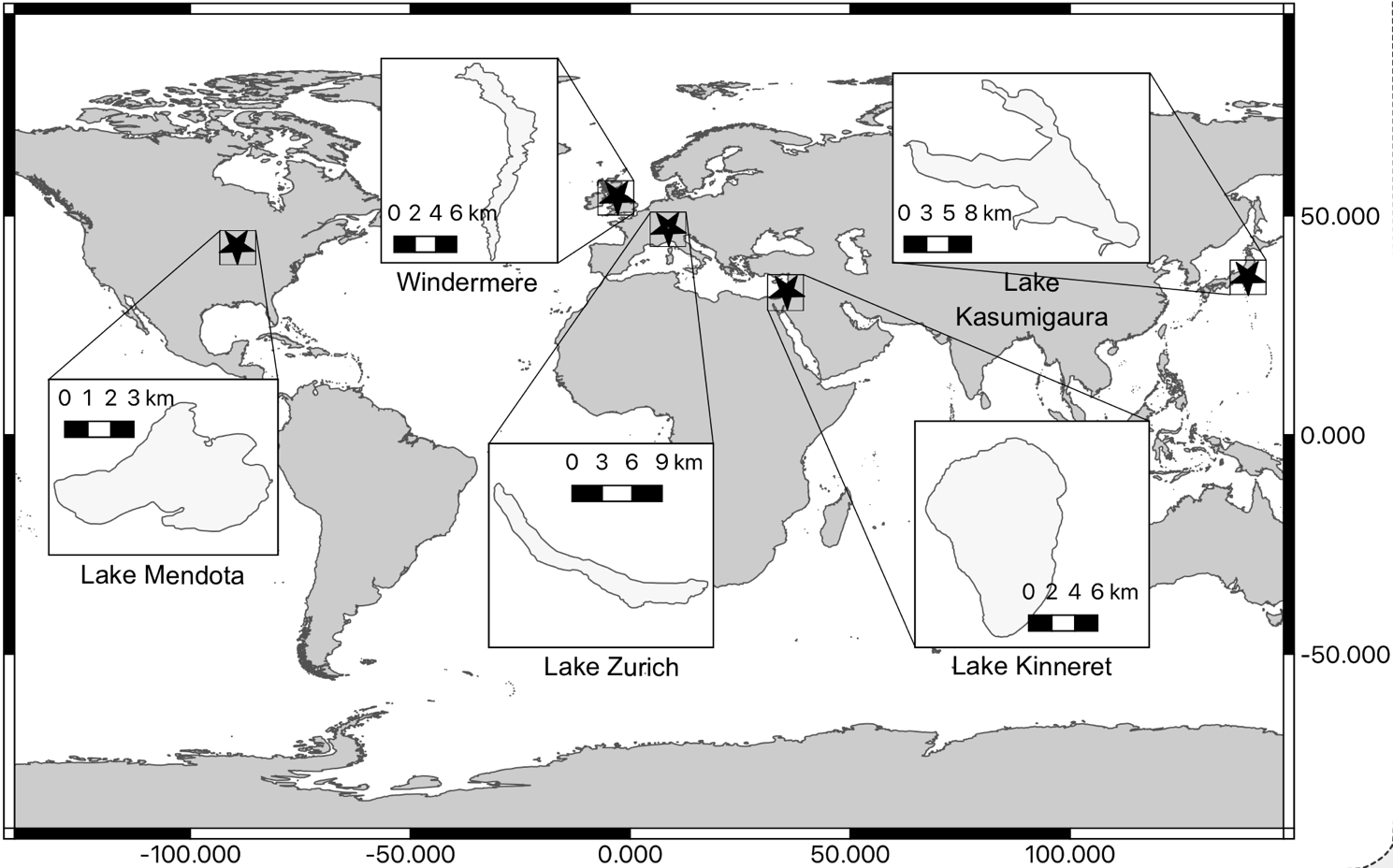

Figure S1. Locations of the five lake ecosystems used in this study and their relative surface areas. Map is projected in WGS 84.

Figure S2. Diagrammatic representation of the three functional diversity metrics based upon trait dissimilarity. Points denote a species’ trait value within the trait space with the size of the point proportional to the abundance of that species at time *t*. Functional richness (*FRic*) quantifies the trait space area encompassed by all species, functional evenness (*FEve*) quantifies the regularity of species within this space and functional dispersion (*FDis*) captures the spread of species relative to the community’s centroid (the ‘centre of gravity’ of all species modulated by abundance).

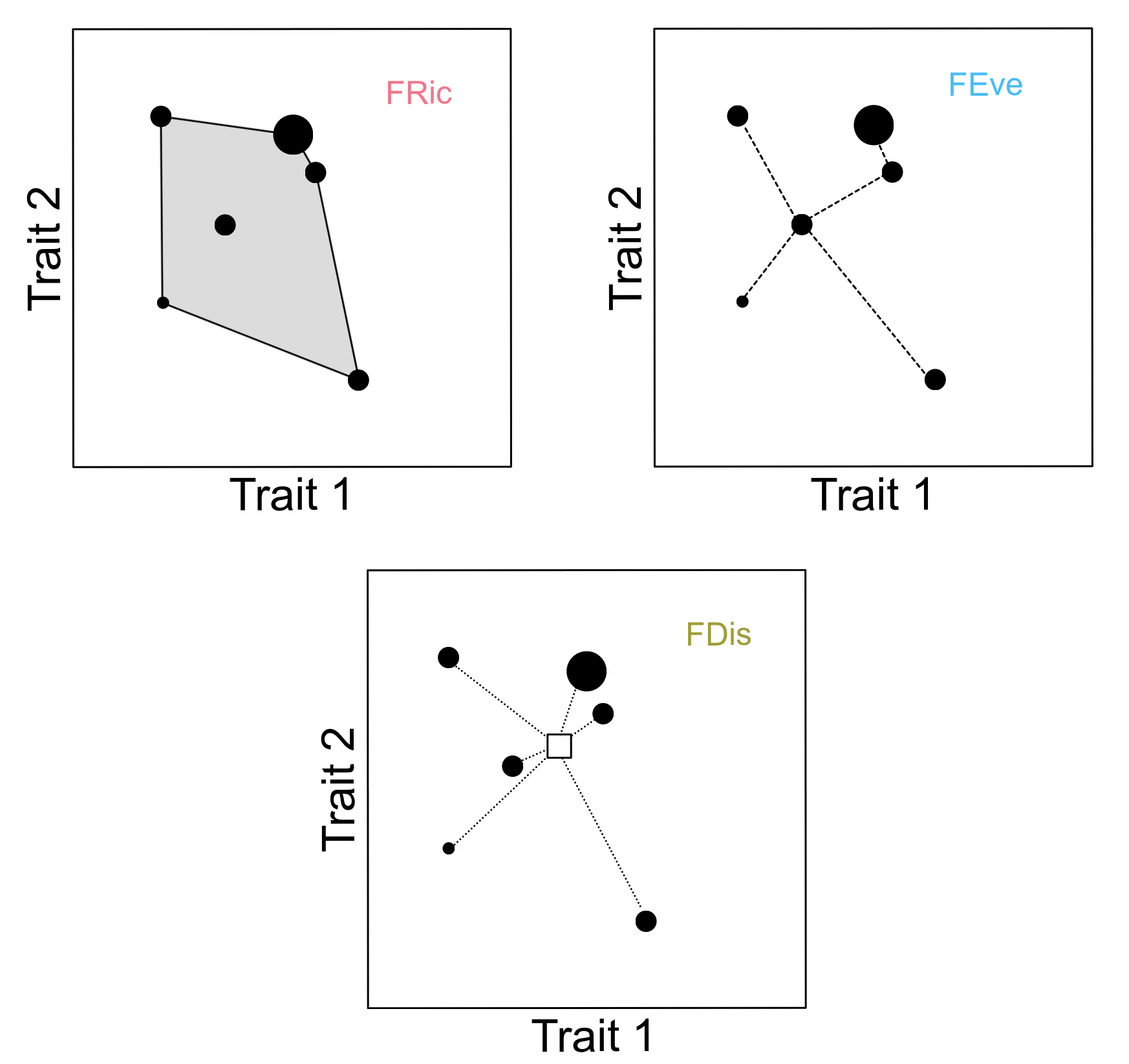

Figure S3. Density of the permuted instantaneous (Lag0) cross-correlation coefficients following 10,000 simulations between functional diversity and each of the system state metrics. Points represent the observed correlation coefficient and stars represent the ‘significant’ relationships (where the observed value transgresses the 2.5-97.5% confidence intervals).

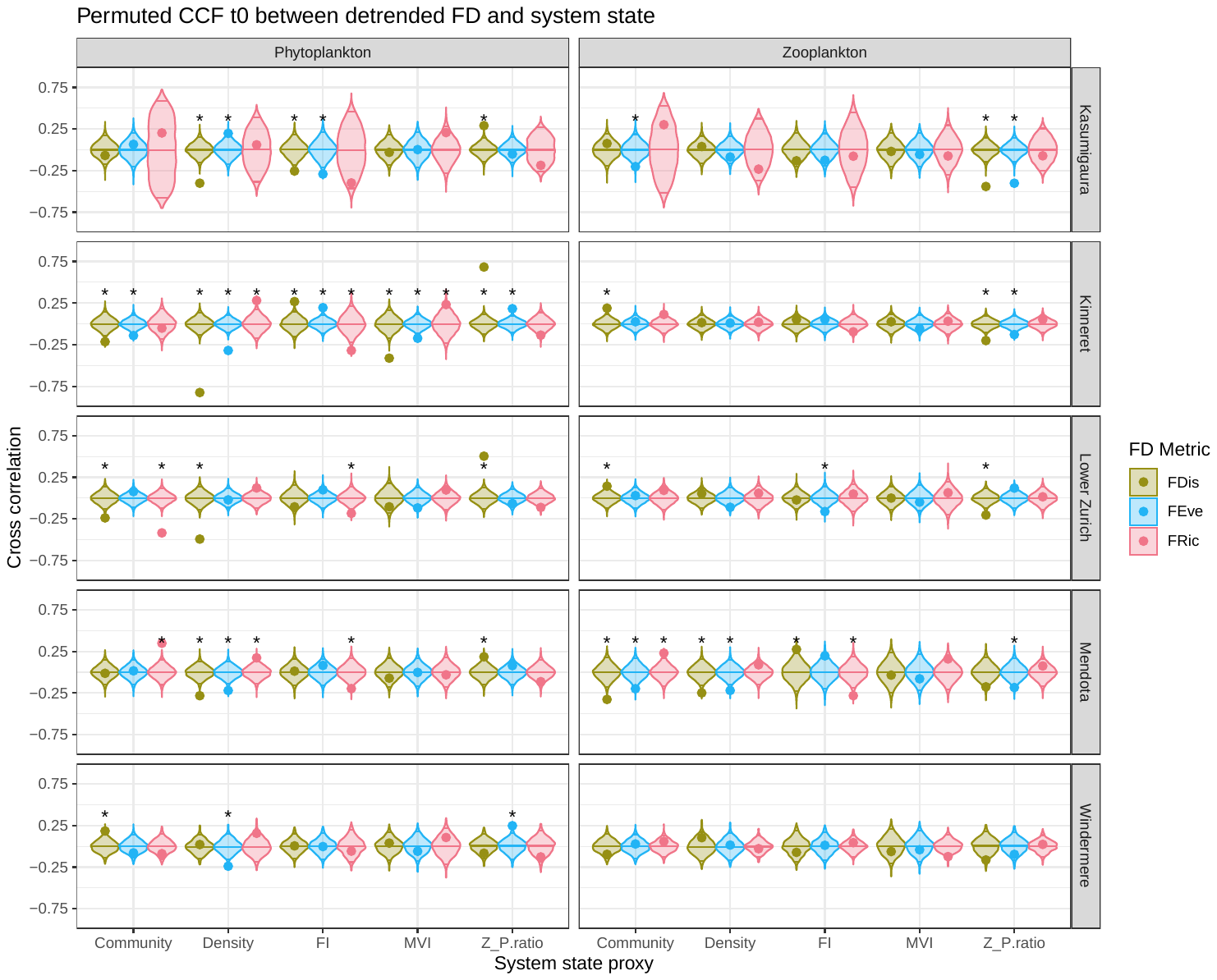

Figure S4. Density of permuted cross-correlation coefficients following 10,000 simulations between functional diversity and each of the system state metrics (LagX). Points represent the absolute strongest observed correlation coefficient and stars represent the ‘significant’ relationships (where the observed value transgresses the 2.5-97.5% confidence intervals). The reported number is the optimal lag (in months) that the observed value was identified. Positive lags indicate functional diversity lagging state, whereas negative lags indicate functional diversity leading state.

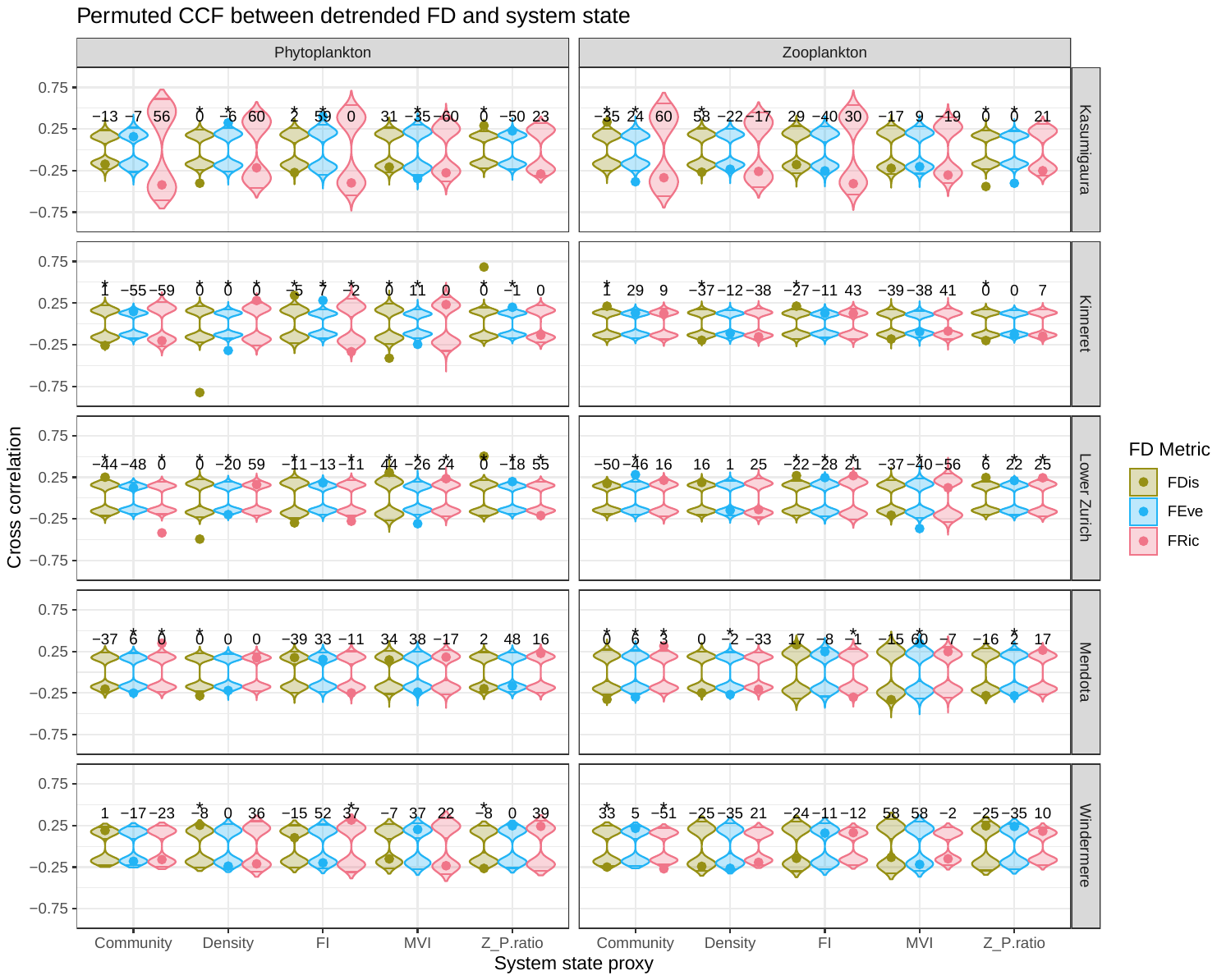

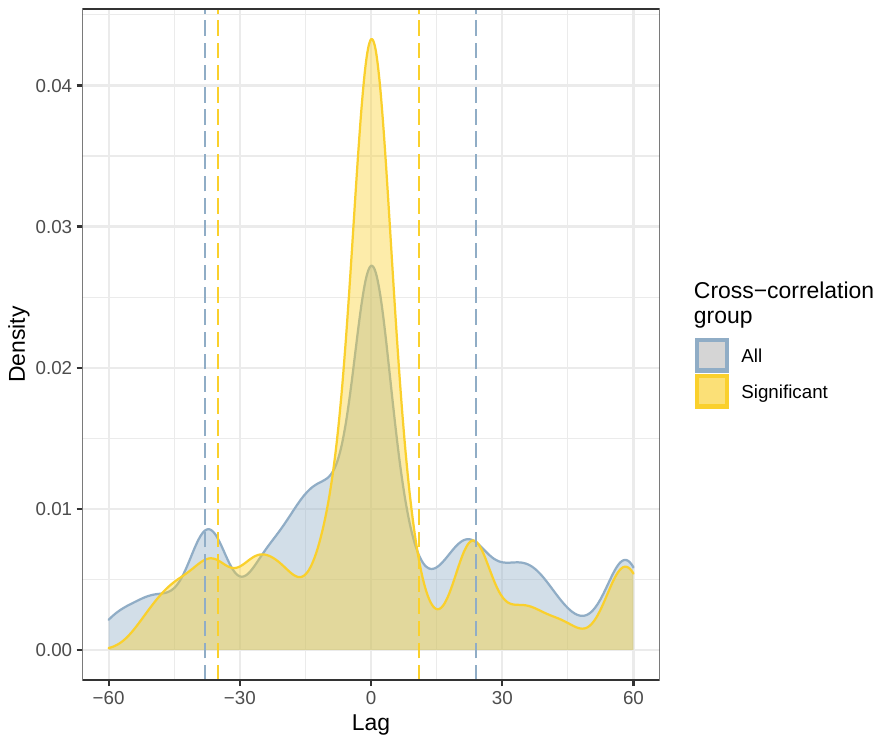

Figure S5. Density plots of the optimal lags identified by cross-correlations pooled across all functional diversity:system state combinations (blue area) or subset to significant combinations only (yellow area). Dashed, vertical lines indicate 10^th^ and 90^th^ quartiles.

Figure S6. Density of instantaneous (Lag0) permuted cross mapping skills where the system state metrics map functional diversity following 1000 simulations. Points represent the observed cross map skill and stars represent the ‘significant’ relationships (where the observed value exceeds the 95% quantile). These points therefore represent the predicted strength of ‘forward’ causality where functional diversity ‘causes’ system state.

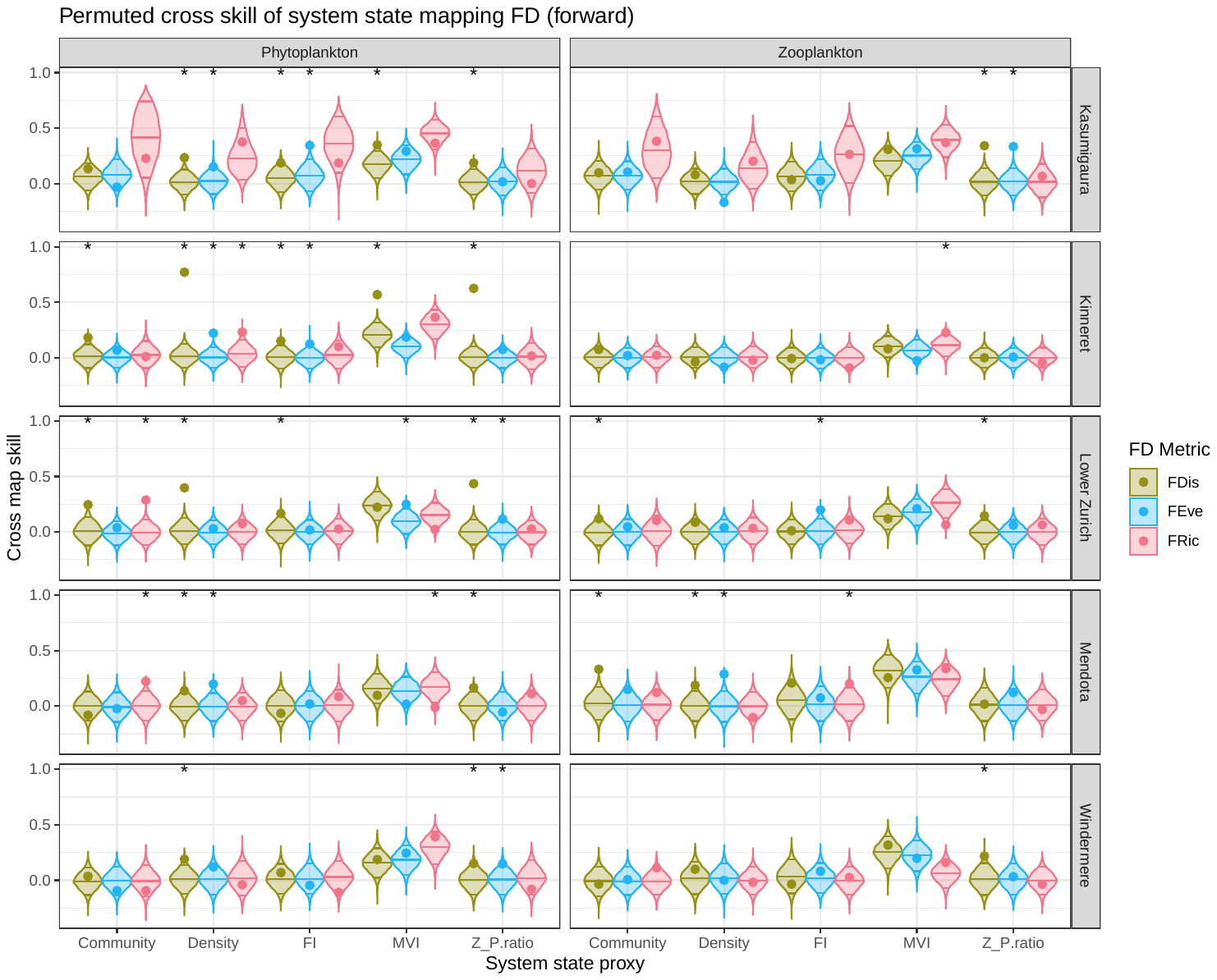

Figure S7. Density of instantaneous (Lag0) permuted cross mapping skills where functional diversity maps system state following 1000 simulations. Points represent the observed cross map skill and stars represent the ‘significant’ relationships (where the observed value exceeds the 95% quantile). These points therefore represent the predicted strength of ‘reverse’ causality where system state ‘causes’ functional diversity.

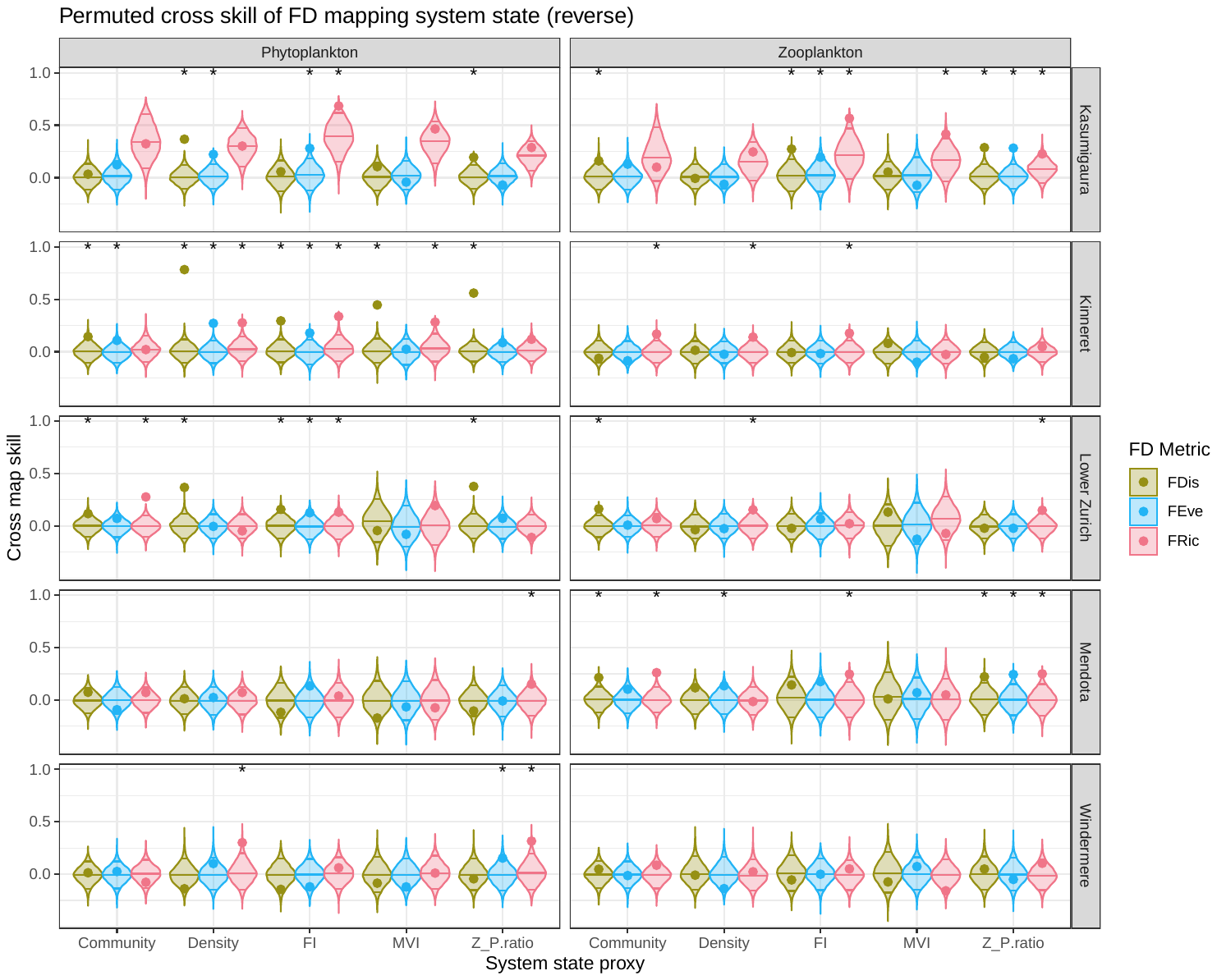

Figure S8. Density of permuted cross mapping skills (LagX) where the system state metrics map functional diversity following 10,000 simulations. Points represent the observed cross map skill and stars represent the ‘significant’ relationships (where the observed value exceeds the 95% quantile). These points therefore represent the predicted strength of ‘forward’ causality where functional diversity ‘causes’ system state. Optimal lags (in months) are indicated by the numbers above each distribution. Negative lags indicate functional diversity leading state whereas positive lags indicate generalised synchrony.

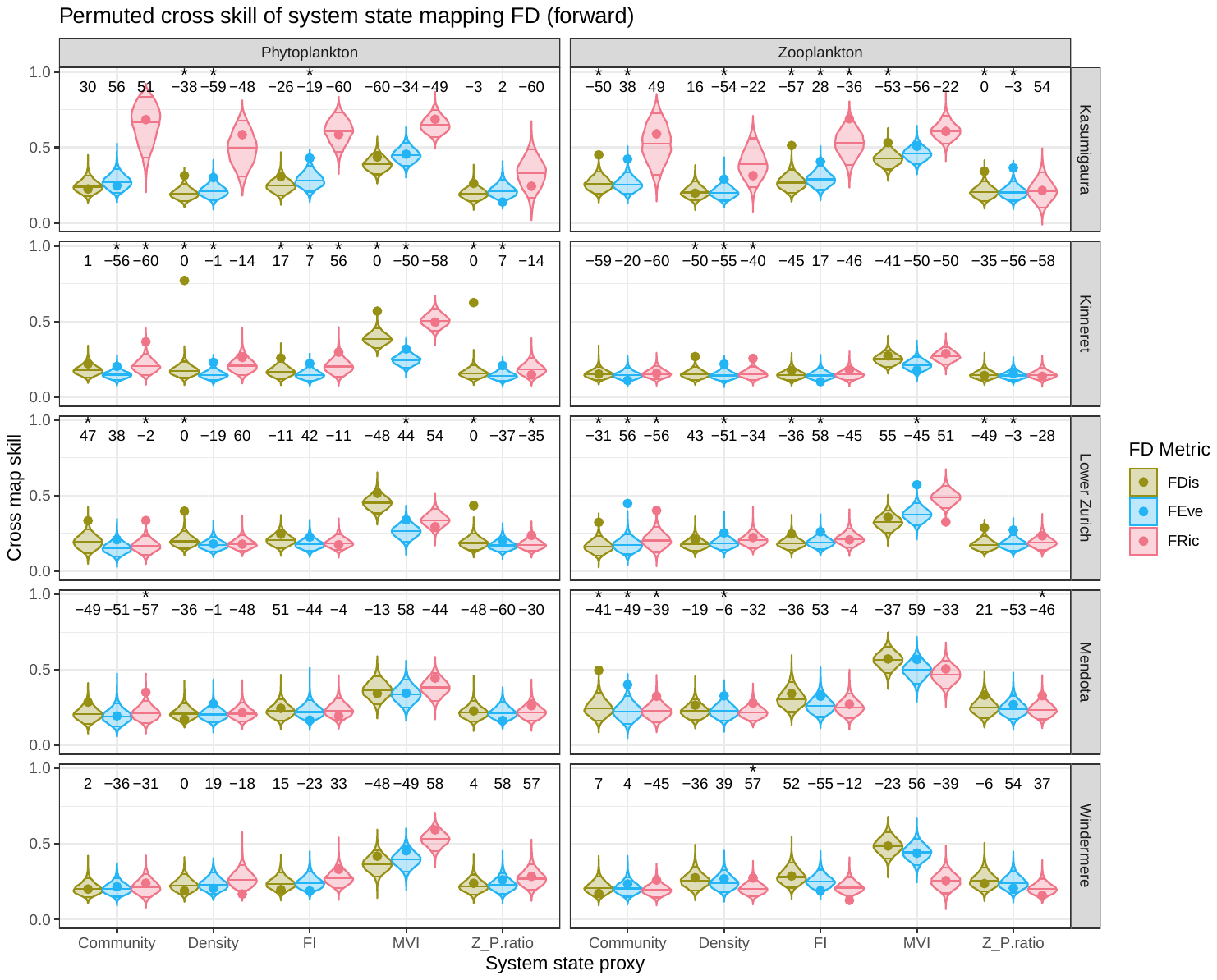

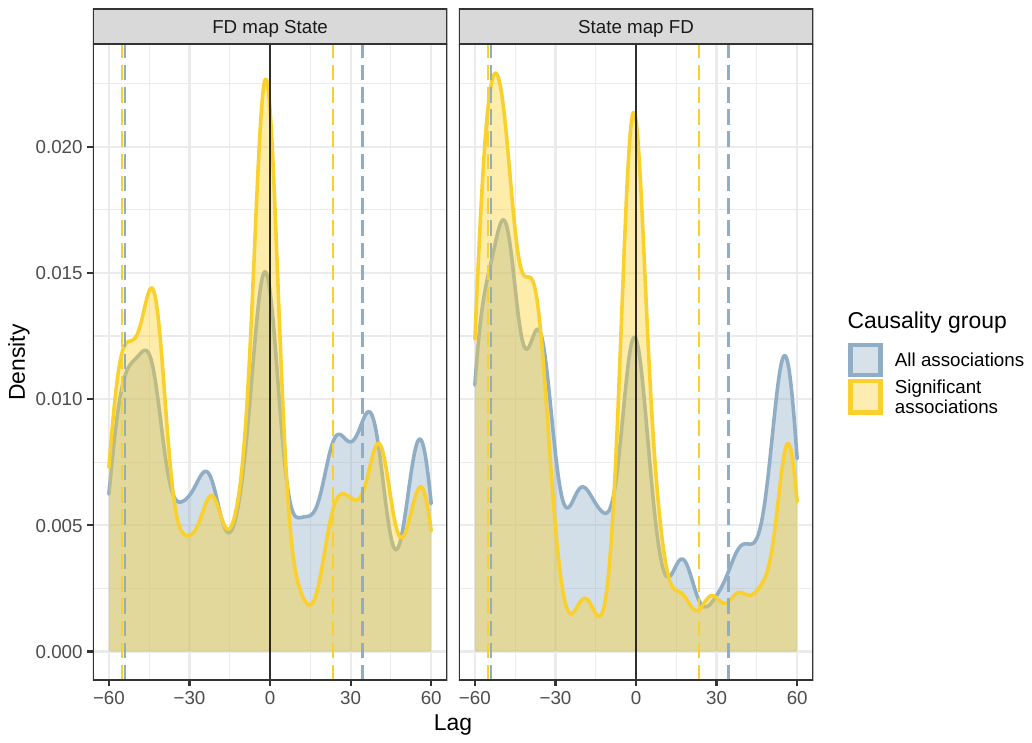

Figure S9. Density plots of the optimal lags identified by forward (State map FD) and reverse (FD map State) cross mappings pooled across all functional diversity:system state combinations (blue area) or subset to significant combinations only (yellow area). Dashed, vertical lines indicate 10^th^ and 90^th^ quartiles.

Figure S10. Density of permuted cross mapping skills (LagX) where functional diversity maps system state following 10,000 simulations. Points represent the observed cross map skill and stars represent the ‘significant’ relationships (where the observed value exceeds the 95% quantile). These points therefore represent the predicted strength of ‘reverse’ causality where system state ‘causes’ functional diversity. Optimal lags (in months) are indicated by the numbers above each distribution. Negative lags indicate functional diversity leading state whereas positive lags indicate generalised synchrony.

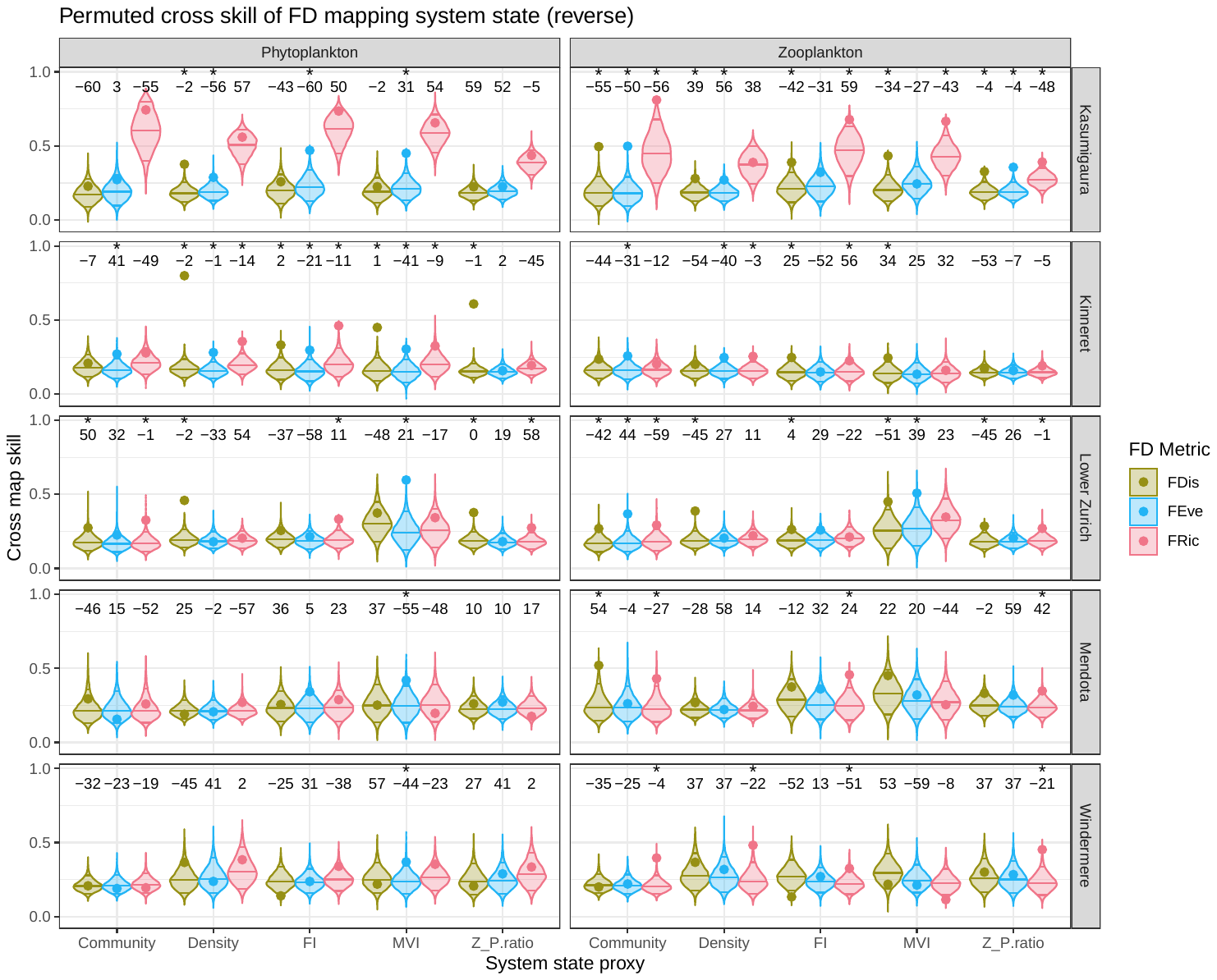

Supplementary Tables

Table S1. Summary statistics for each functional diversity-system state correlation at Lag0. Variation is reported as standard errors.

| **Trophic Guild** | **Functional diversity metric** | **System state metric** | **Median correlation** | **Number of significant lakes** | **Proportion significant** |
| --- | --- | --- | --- | --- | --- |
| Phytoplankton | FDis | Community | -0.068 ± 0.034 | 3 | 0.6 |
|  |  | Density | -0.402 ± 0.062 | 4 | 0.8 |
|  |  | FI | 0.005 ± 0.038 | 2 | 0.4 |
|  |  | MVI | -0.072 ± 0.035 | 1 | 0.2 |
|  |  | Z_P.ratio | 0.29 ± 0.059 | 4 | 0.8 |
|  | FEve | Community | 0.016 ± 0.019 | 1 | 0.2 |
|  |  | Density | -0.22 ± 0.042 | 4 | 0.8 |
|  |  | FI | 0.081 ± 0.037 | 2 | 0.4 |
|  |  | MVI | -0.059 ± 0.015 | 1 | 0.2 |
|  |  | Z_P.ratio | 0.079 ± 0.028 | 2 | 0.4 |
|  | FRic | Community | -0.051 ± 0.059 | 2 | 0.4 |
|  |  | Density | 0.157 ± 0.016 | 2 | 0.4 |
|  |  | FI | -0.197 ± 0.026 | 3 | 0.6 |
|  |  | MVI | 0.106 ± 0.021 | 1 | 0.2 |
|  |  | Z_P.ratio | -0.127 ± 0.006 | 0 | 0 |
| Zooplankton | FDis | Community | 0.075 ± 0.042 | 3 | 0.6 |
|  |  | Density | 0.039 ± 0.028 | 1 | 0.2 |
|  |  | FI | -0.023 ± 0.032 | 1 | 0.2 |
|  |  | MVI | -0.02 ± 0.007 | 0 | 0 |
|  |  | Z_P.ratio | -0.2 ± 0.023 | 3 | 0.6 |
|  | FEve | Community | 0.026 ± 0.025 | 2 | 0.4 |
|  |  | Density | -0.087 ± 0.019 | 1 | 0.2 |
|  |  | FI | 0.012 ± 0.029 | 1 | 0.2 |
|  |  | MVI | -0.056 ± 0.003 | 0 | 0 |
|  |  | Z_P.ratio | -0.128 ± 0.037 | 3 | 0.6 |
|  | FRic | Community | 0.114 ± 0.02 | 1 | 0.2 |
|  |  | Density | 0.02 ± 0.026 | 0 | 0 |
|  |  | FI | -0.077 ± 0.027 | 1 | 0.2 |
|  |  | MVI | 0.032 ± 0.022 | 0 | 0 |
|  |  | Z_P.ratio | 0.023 ± 0.011 | 0 | 0 |

| **Trophic Guild** | **Functional diversity metric** | **System state metric** | **Median correlation** | **Median lag** | **Number of significant lakes** | **Proportion significant** |
| --- | --- | --- | --- | --- | --- | --- |
| Phytoplankton | FDis | Community | -0.173 ± 0.048 | -13 ± 4.22 | 2 | 0.4 |
|  |  | Density | -0.402 ± 0.078 | 0 ± 0.72 | 5 | 1 |
|  |  | FI | 0.104 ± 0.057 | -11 ± 3.12 | 3 | 0.6 |
|  |  | MVI | -0.15 ± 0.058 | 31 ± 4.5 | 2 | 0.4 |
|  |  | Z_P.ratio | 0.29 ± 0.084 | 0 ± 0.78 | 4 | 0.8 |
|  | FEve | Community | 0.124 ± 0.04 | -17 ± 5.27 | 1 | 0.2 |
|  |  | Density | -0.22 ± 0.052 | 0 ± 1.73 | 3 | 0.6 |
|  |  | FI | 0.183 ± 0.045 | 33 ± 6.07 | 2 | 0.4 |
|  |  | MVI | -0.246 ± 0.045 | 11 ± 6.86 | 3 | 0.6 |
|  |  | Z_P.ratio | 0.2 ± 0.035 | -1 ± 7.1 | 2 | 0.4 |
|  | FRic | Community | -0.203 ± 0.063 | 0 ± 8.37 | 2 | 0.4 |
|  |  | Density | 0.162 ± 0.047 | 36 ± 5.98 | 1 | 0.2 |
|  |  | FI | -0.278 ± 0.058 | -2 ± 3.98 | 3 | 0.6 |
|  |  | MVI | 0.181 ± 0.052 | 0 ± 6.9 | 1 | 0.2 |
|  |  | Z_P.ratio | -0.135 ± 0.05 | 23 ± 4.23 | 1 | 0.2 |
| Zooplankton | FDis | Community | 0.174 ± 0.059 | 0 ± 6.55 | 4 | 0.8 |
|  |  | Density | -0.241 ± 0.038 | 0 ± 7.47 | 2 | 0.4 |
|  |  | FI | 0.213 ± 0.048 | -22 ± 5.27 | 2 | 0.4 |
|  |  | MVI | -0.206 ± 0.015 | -17 ± 7.92 | 0 | 0 |
|  |  | Z_P.ratio | -0.2 ± 0.063 | 0 ± 2.59 | 3 | 0.6 |
|  | FEve | Community | 0.131 ± 0.062 | 6 ± 5.94 | 3 | 0.6 |
|  |  | Density | -0.234 ± 0.014 | -12 ± 2.96 | 1 | 0.2 |
|  |  | FI | 0.159 ± 0.041 | -11 ± 2.77 | 1 | 0.2 |
|  |  | MVI | -0.203 ± 0.054 | 9 ± 9.8 | 2 | 0.4 |
|  |  | Z_P.ratio | -0.128 ± 0.058 | 0 ± 4.11 | 3 | 0.6 |
|  | FRic | Community | 0.125 ± 0.059 | 9 ± 7.92 | 2 | 0.4 |
|  |  | Density | -0.192 ± 0.01 | -17 ± 5.95 | 0 | 0 |
|  |  | FI | 0.126 ± 0.061 | 21 ± 4.5 | 2 | 0.4 |
|  |  | MVI | -0.086 ± 0.044 | -7 ± 6.97 | 0 | 0 |
|  |  | Z_P.ratio | 0.182 ± 0.048 | 17 ± 1.5 | 1 | 0.2 |

Table S2. Summary statistics for each functional diversity-system state cross correlation across lags (LagX). Variation is reported as standard errors.

Table S3. Summary statistics for each functional diversity-system state cross mapping at Lag0. This is the forward relationship (system state maps functional diversity) where a significant relationship suggests diversity causes state. Variation is reported as standard errors.

| **Trophic Guild** | **Functional diversity metric** | **System state metric** | **Median correlation** | **Number of significant lakes** | **Proportion significant** |
| --- | --- | --- | --- | --- | --- |
| Phytoplankton | FDis | Community | 0.134 ± 0.03 | 2 | 0.4 |
|  |  | Density | 0.233 ± 0.05 | 5 | 1 |
|  |  | FI | 0.152 ± 0.02 | 3 | 0.6 |
|  |  | MVI | 0.223 ± 0.04 | 2 | 0.4 |
|  |  | Z_P.ratio | 0.188 ± 0.04 | 5 | 1 |
|  | FEve | Community | -0.023 ± 0.01 | 0 | 0 |
|  |  | Density | 0.152 ± 0.02 | 3 | 0.6 |
|  |  | FI | 0.018 ± 0.03 | 2 | 0.4 |
|  |  | MVI | 0.244 ± 0.02 | 1 | 0.2 |
|  |  | Z_P.ratio | 0.075 ± 0.02 | 2 | 0.4 |
|  | FRic | Community | 0.222 ± 0.03 | 2 | 0.4 |
|  |  | Density | 0.074 ± 0.03 | 1 | 0.2 |
|  |  | FI | 0.084 ± 0.02 | 0 | 0 |
|  |  | MVI | 0.364 ± 0.04 | 1 | 0.2 |
|  |  | Z_P.ratio | 0.017 ± 0.01 | 0 | 0 |
| Zooplankton | FDis | Community | 0.098 ± 0.03 | 2 | 0.4 |
|  |  | Density | 0.087 ± 0.02 | 1 | 0.2 |
|  |  | FI | 0.01 ± 0.02 | 0 | 0 |
|  |  | MVI | 0.255 ± 0.02 | 0 | 0 |
|  |  | Z_P.ratio | 0.144 ± 0.03 | 3 | 0.6 |
|  | FEve | Community | 0.046 ± 0.01 | 0 | 0 |
|  |  | Density | 0 ± 0.04 | 1 | 0.2 |
|  |  | FI | 0.074 ± 0.02 | 1 | 0.2 |
|  |  | MVI | 0.209 ± 0.03 | 0 | 0 |
|  |  | Z_P.ratio | 0.06 ± 0.03 | 1 | 0.2 |
|  | FRic | Community | 0.111 ± 0.03 | 0 | 0 |
|  |  | Density | -0.019 ± 0.02 | 0 | 0 |
|  |  | FI | 0.107 ± 0.03 | 1 | 0.2 |
|  |  | MVI | 0.227 ± 0.03 | 1 | 0.2 |
|  |  | Z_P.ratio | -0.032 ± 0.01 | 0 | 0 |

Table S4. Summary statistics for each functional diversity-system state cross mapping at Lag0. This is the reverse relationship (functional diversity maps system state) where a significant relationship suggests diversity is caused by state. Variation is reported as standard errors.

| **Trophic Guild** | **Functional diversity metric** | **System state metric** | **Median correlation** | **Number of significant lakes** | **Proportion significant** |
| --- | --- | --- | --- | --- | --- |
| Phytoplankton | FDis | Community | 0.074 ± 0.01 | 2 | 0.4 |
|  |  | Density | 0.366 ± 0.07 | 3 | 0.6 |
|  |  | FI | 0.057 ± 0.04 | 2 | 0.4 |
|  |  | MVI | -0.045 ± 0.05 | 1 | 0.2 |
|  |  | Z_P.ratio | 0.194 ± 0.06 | 3 | 0.6 |
|  | FEve | Community | 0.071 ± 0.02 | 1 | 0.2 |
|  |  | Density | 0.102 ± 0.02 | 2 | 0.4 |
|  |  | FI | 0.134 ± 0.03 | 3 | 0.6 |
|  |  | MVI | -0.065 ± 0.01 | 0 | 0 |
|  |  | Z_P.ratio | 0.072 ± 0.02 | 1 | 0.2 |
|  | FRic | Community | 0.072 ± 0.03 | 1 | 0.2 |
|  |  | Density | 0.276 ± 0.03 | 2 | 0.4 |
|  |  | FI | 0.131 ± 0.05 | 3 | 0.6 |
|  |  | MVI | 0.194 ± 0.04 | 1 | 0.2 |
|  |  | Z_P.ratio | 0.152 ± 0.03 | 2 | 0.4 |
| Zooplankton | FDis | Community | 0.16 ± 0.02 | 3 | 0.6 |
|  |  | Density | -0.007 ± 0.01 | 0 | 0 |
|  |  | FI | -0.008 ± 0.03 | 1 | 0.2 |
|  |  | MVI | 0.053 ± 0.02 | 0 | 0 |
|  |  | Z_P.ratio | 0.05 ± 0.03 | 2 | 0.4 |
|  | FEve | Community | 0.007 ± 0.02 | 0 | 0 |
|  |  | Density | -0.026 ± 0.02 | 1 | 0.2 |
|  |  | FI | 0.066 ± 0.02 | 1 | 0.2 |
|  |  | MVI | -0.073 ± 0.02 | 0 | 0 |
|  |  | Z_P.ratio | -0.022 ± 0.03 | 2 | 0.4 |
|  | FRic | Community | 0.099 ± 0.02 | 2 | 0.4 |
|  |  | Density | 0.14 ± 0.02 | 2 | 0.4 |
|  |  | FI | 0.177 ± 0.04 | 3 | 0.6 |
|  |  | MVI | -0.026 ± 0.04 | 1 | 0.2 |
|  |  | Z_P.ratio | 0.15 ± 0.02 | 3 | 0.6 |

Table S5. Causality comparison for lag0/instantaneous cross mappings between functional diversity and system state. The proportion of lakes with forward, reverse, bi-directional and zero causality are reported. Variation is reported as standard errors.

| **Trophic Guild** | **Functional diversity metric** | **System state metric** | **Proportion of lakes forward causal** | **Proportion of lakes reverse causal** | **Proportion of lakes bi-directionally causal** | **Proportion of lakes non-causal** |
| --- | --- | --- | --- | --- | --- | --- |
| Phytoplankton | FDis | Community | 0 | 0 | 0.4 | 0.6 |
|  |  | Density | 0.4 | 0 | 0.6 | 0 |
|  |  | FI | 0.2 | 0 | 0.4 | 0.4 |
|  |  | MVI | 0.2 | 0 | 0.2 | 0.6 |
|  |  | Z_P.ratio | 0.4 | 0 | 0.6 | 0 |
|  | FEve | Community | 0 | 0.2 | 0 | 0.8 |
|  |  | Density | 0.2 | 0 | 0.4 | 0.4 |
|  |  | FI | 0 | 0.2 | 0.4 | 0.4 |
|  |  | MVI | 0.2 | 0 | 0 | 0.8 |
|  |  | Z_P.ratio | 0.2 | 0 | 0.2 | 0.6 |
|  | FRic | Community | 0.2 | 0 | 0.2 | 0.6 |
|  |  | Density | 0 | 0.2 | 0.2 | 0.6 |
|  |  | FI | 0 | 0.6 | 0 | 0.4 |
|  |  | MVI | 0.2 | 0.2 | 0 | 0.6 |
|  |  | Z_P.ratio | 0 | 0.4 | 0 | 0.6 |
| Zooplankton | FDis | Community | 0 | 0.2 | 0.4 | 0.4 |
|  |  | Density | 0.2 | 0 | 0 | 0.8 |
|  |  | FI | 0 | 0.2 | 0 | 0.8 |
|  |  | MVI | 0 | 0 | 0 | 1 |
|  |  | Z_P.ratio | 0.4 | 0.2 | 0.2 | 0.2 |
|  | FEve | Community | 0 | 0 | 0 | 1 |
|  |  | Density | 0 | 0 | 0.2 | 0.8 |
|  |  | FI | 0.2 | 0.2 | 0 | 0.6 |
|  |  | MVI | 0 | 0 | 0 | 1 |
|  |  | Z_P.ratio | 0 | 0.2 | 0.2 | 0.6 |
|  | FRic | Community | 0 | 0.4 | 0 | 0.6 |
|  |  | Density | 0 | 0.4 | 0 | 0.6 |
|  |  | FI | 0 | 0.4 | 0.2 | 0.4 |
|  |  | MVI | 0.2 | 0.2 | 0 | 0.6 |
|  |  | Z_P.ratio | 0 | 0.6 | 0 | 0.4 |

Table S6. Summary statistics for each functional diversity-system state cross mapping across lags (LagX). This is the forward relationship (system state maps functional diversity) where a significant relationship suggests diversity causes state. Variation is reported as standard errors.

| **Trophic Guild** | **Functional diversity metric** | **System state metric** | **Median correlation** | **Median lag** | **Number of significant lakes** | **Proportion significant** |
| --- | --- | --- | --- | --- | --- | --- |
| Phytoplankton | FDis | Community | 0.224 ± 0.01 | 2 ± 7.30 | 1 | 0.2 |
|  |  | Density | 0.314 ± 0.05 | 0 ± 4.06 | 3 | 0.6 |
|  |  | FI | 0.245 ± 0.01 | 15 ± 5.91 | 1 | 0.2 |
|  |  | MVI | 0.436 ± 0.02 | -48 ± 5.16 | 1 | 0.2 |
|  |  | Z_P.ratio | 0.262 ± 0.03 | 0 ± 4.34 | 2 | 0.4 |
|  | FEve | Community | 0.209 ± 0.00 | -36 ± 10.55 | 1 | 0.2 |
|  |  | Density | 0.23 ± 0.01 | -1 ± 5.88 | 2 | 0.4 |
|  |  | FI | 0.221 ± 0.02 | -19 ± 6.61 | 2 | 0.4 |
|  |  | MVI | 0.345 ± 0.01 | -34 ± 10.57 | 2 | 0.4 |
|  |  | Z_P.ratio | 0.202 ± 0.01 | 2 ± 9.06 | 1 | 0.2 |
|  | FRic | Community | 0.352 ± 0.03 | -31 ± 9.20 | 3 | 0.6 |
|  |  | Density | 0.216 ± 0.04 | -18 ± 8.83 | 0 | 0 |
|  |  | FI | 0.297 ± 0.03 | -4 ± 8.90 | 1 | 0.2 |
|  |  | MVI | 0.496 ± 0.03 | -44 ± 11.70 | 0 | 0 |
|  |  | Z_P.ratio | 0.243 ± 0.01 | -30 ± 8.85 | 1 | 0.2 |
| Zooplankton | FDis | Community | 0.324 ± 0.03 | -41 ± 5.12 | 3 | 0.6 |
|  |  | Density | 0.266 ± 0.01 | -19 ± 7.64 | 1 | 0.2 |
|  |  | FI | 0.288 ± 0.03 | -36 ± 8.71 | 2 | 0.4 |
|  |  | MVI | 0.486 ± 0.03 | -37 ± 8.63 | 1 | 0.2 |
|  |  | Z_P.ratio | 0.289 ± 0.02 | -6 ± 5.61 | 2 | 0.4 |
|  | FEve | Community | 0.402 ± 0.03 | 4 ± 8.50 | 3 | 0.6 |
|  |  | Density | 0.269 ± 0.01 | -51 ± 8.29 | 4 | 0.8 |
|  |  | FI | 0.261 ± 0.02 | 28 ± 9.07 | 2 | 0.4 |
|  |  | MVI | 0.507 ± 0.03 | -45 ± 11.84 | 1 | 0.2 |
|  |  | Z_P.ratio | 0.27 ± 0.02 | -3 ± 9.02 | 2 | 0.4 |
|  | FRic | Community | 0.325 ± 0.03 | -45 ± 9.01 | 2 | 0.4 |
|  |  | Density | 0.275 ± 0.01 | -32 ± 8.07 | 2 | 0.4 |
|  |  | FI | 0.208 ± 0.05 | -36 ± 3.88 | 1 | 0.2 |
|  |  | MVI | 0.326 ± 0.03 | -33 ± 8.04 | 0 | 0 |
|  |  | Z_P.ratio | 0.215 ± 0.02 | -28 ± 10.11 | 1 | 0.2 |

| **Trophic Guild** | **Functional diversity metric** | **System state metric** | **Median correlation** | **Median lag** | **Number of significant lakes** | **Proportion significant** |
| --- | --- | --- | --- | --- | --- | --- |
| Phytoplankton | FDis | Community | 0.228 ± 0.008 | -32 ± 8.65 | 1 | 0.2 |
|  |  | Density | 0.376 ± 0.045 | -2 ± 5.03 | 3 | 0.6 |
|  |  | FI | 0.256 ± 0.014 | -25 ± 6.52 | 1 | 0.2 |
|  |  | MVI | 0.252 ± 0.021 | 1 ± 8.08 | 1 | 0.2 |
|  |  | Z_P.ratio | 0.261 ± 0.033 | 10 ± 5.01 | 2 | 0.4 |
|  | FEve | Community | 0.225 ± 0.01 | 15 ± 5.04 | 1 | 0.2 |
|  |  | Density | 0.238 ± 0.009 | -2 ± 7.34 | 2 | 0.4 |
|  |  | FI | 0.297 ± 0.02 | -21 ± 7.92 | 2 | 0.4 |
|  |  | MVI | 0.42 ± 0.022 | -41 ± 8.06 | 5 | 1 |
|  |  | Z_P.ratio | 0.226 ± 0.011 | 19 ± 4.21 | 0 | 0 |
|  | FRic | Community | 0.279 ± 0.044 | -49 ± 4.79 | 1 | 0.2 |
|  |  | Density | 0.356 ± 0.027 | 2 ± 9.62 | 1 | 0.2 |
|  |  | FI | 0.339 ± 0.036 | 11 ± 6.69 | 2 | 0.4 |
|  |  | MVI | 0.342 ± 0.034 | -17 ± 7.58 | 1 | 0.2 |
|  |  | Z_P.ratio | 0.274 ± 0.021 | 2 ± 7.46 | 1 | 0.2 |
| Zooplankton | FDis | Community | 0.268 ± 0.03 | -42 ± 8.88 | 3 | 0.6 |
|  |  | Density | 0.28 ± 0.015 | -28 ± 9 | 2 | 0.4 |
|  |  | FI | 0.263 ± 0.021 | -12 ± 6.38 | 3 | 0.6 |
|  |  | MVI | 0.433 ± 0.023 | 22 ± 8.99 | 3 | 0.6 |
|  |  | Z_P.ratio | 0.3 ± 0.013 | -4 ± 7.3 | 2 | 0.4 |
|  | FEve | Community | 0.262 ± 0.023 | -25 ± 7.19 | 3 | 0.6 |
|  |  | Density | 0.248 ± 0.009 | 37 ± 7.99 | 2 | 0.4 |
|  |  | FI | 0.27 ± 0.016 | 13 ± 7.54 | 1 | 0.2 |
|  |  | MVI | 0.244 ± 0.028 | 20 ± 8.22 | 1 | 0.2 |
|  |  | Z_P.ratio | 0.284 ± 0.016 | 26 ± 5.59 | 1 | 0.2 |
|  | FRic | Community | 0.396 ± 0.046 | -27 ± 5.01 | 4 | 0.8 |
|  |  | Density | 0.255 ± 0.023 | 11 ± 4.43 | 2 | 0.4 |
|  |  | FI | 0.325 ± 0.039 | 24 ± 9.7 | 3 | 0.6 |
|  |  | MVI | 0.255 ± 0.044 | -8 ± 7.13 | 1 | 0.2 |
|  |  | Z_P.ratio | 0.348 ± 0.02 | -5 ± 6.57 | 4 | 0.8 |

Table S7. Summary statistics for each functional diversity-system state cross mapping across lags (LagX). This is the reverse relationship (functional diversity maps system state) where a significant relationship suggests diversity is caused by state. Variation is reported as standard errors.

| **Trophic Guild**  Table S8. Causality comparison for cross mappings between functional diversity and system state, across lags. The proportion of lakes with forward, reverse, bi-directional and zero causality are reported. Variation is reported as standard errors. | **Functional diversity metric** | **System state metric** | **Proportion of lakes forward causal** | **Proportion of lakes reverse causal** | **Proportion of lakes bi-directionally causal** | **Proportion of lakes non-causal** | **Median difference in lag (lagReverse – lagForward)** |
| --- | --- | --- | --- | --- | --- | --- | --- |
| Phytoplankton | FDis | Community | 0 | 0 | 0.2 | 0.8 | -25.2 |
|  |  | Density | 0 | 0 | 0.6 | 0.4 | 9.6 |
|  |  | FI | 0 | 0 | 0.2 | 0.8 | -22.6 |
|  |  | MVI | 0 | 0 | 0.2 | 0.8 | 42.8 |
|  |  | Z_P.ratio | 0 | 0 | 0.4 | 0.6 | 28.4 |
|  | FEve | Community | 0 | 0 | 0.2 | 0.8 | 23.4 |
|  |  | Density | 0 | 0 | 0.4 | 0.6 | 2 |
|  |  | FI | 0 | 0 | 0.4 | 0.6 | -13.2 |
|  |  | MVI | 0 | 0.6 | 0.4 | 0 | -11.4 |
|  |  | Z_P.ratio | 0.2 | 0 | 0 | 0.8 | 30.8 |
|  | FRic | Community | 0.4 | 0 | 0.2 | 0.4 | -15.4 |
|  |  | Density | 0 | 0.2 | 0 | 0.8 | 22 |
|  |  | FI | 0 | 0.2 | 0.2 | 0.6 | 4.2 |
|  |  | MVI | 0 | 0.2 | 0 | 0.8 | -0.8 |
|  |  | Z_P.ratio | 0 | 0 | 0.2 | 0.8 | 21.8 |
| Zooplankton | FDis | Community | 0 | 0 | 0.6 | 0.4 | 10.4 |
|  |  | Density | 0.2 | 0.4 | 0 | 0.4 | -1 |
|  |  | FI | 0 | 0.2 | 0.4 | 0.4 | 9 |
|  |  | MVI | 0 | 0.4 | 0.2 | 0.4 | 24.6 |
|  |  | Z_P.ratio | 0 | 0 | 0.4 | 0.6 | 0.4 |
|  | FEve | Community | 0.2 | 0.2 | 0.4 | 0.2 | -19 |
|  |  | Density | 0.4 | 0 | 0.4 | 0.2 | 53 |
|  |  | FI | 0.4 | 0 | 0 | 0.6 | -22 |
|  |  | MVI | 0 | 0 | 0.2 | 0.8 | 6.8 |
|  |  | Z_P.ratio | 0.2 | 0 | 0.2 | 0.6 | 34.4 |
|  | FRic | Community | 0 | 0.4 | 0.4 | 0.2 | -1.4 |
|  |  | Density | 0 | 0 | 0.4 | 0.6 | 21.8 |
|  |  | FI | 0 | 0.6 | 0.2 | 0.2 | 41.8 |
|  |  | MVI | 0 | 0.2 | 0 | 0.8 | 10.6 |
|  |  | Z_P.ratio | 0 | 0.6 | 0.2 | 0.2 | 1.6 |
